# MerMAIDs: A novel family of metagenomically discovered, marine, anion-conducting and intensely desensitizing channelrhodopsins

**DOI:** 10.1101/607804

**Authors:** Johannes Oppermann, Paul Fischer, Arita Silapetere, Bernhard Liepe, Silvia Rodriguez-Rozada, José Flores-Uribe, Enrico Peter, Anke Keidel, Johannes Vierock, Joel Kaufmann, Matthias Broser, Meike Luck, Franz Bartl, Peter Hildebrandt, J. Simon Wiegert, Oded Béjà, Peter Hegemann, Jonas Wietek

**Author notes:** Corresponding authors. (P.H.); (J.W.).

## Abstract

Channelrhodopsins (ChRs) are algal light-gated ion channels widely used as optogenetic tools for manipulating neuronal activity. ChRs desensitize under continuous bright-light illumination, resulting in a significant decline of photocurrents. We describe a novel, metagenomically identified family of phylogenetically distinct anion-conducting ChRs (designated MerMAIDs). MerMAIDs almost completely desensitize during continuous illumination due to accumulation of a late non-conducting photointermediate that disrupts the ion permeation pathway. MerMAID desensitization can be fully explained by a single photocycle in which a long-lived desensitized state follows the short-lived conducting state. A conserved cysteine is the critical factor in desensitization, as its mutation results in recovery of large stationary photocurrents. The rapid desensitization of MerMAIDs enables their use as optogenetic silencers for transient suppression of individual action potentials without affecting subsequent spiking during continuous illumination. Our results could facilitate the development of further novel optogenetic tools from metagenomic databases and enhance general understanding of ChR function.

---

Channelrhodopsins (ChRs) are members of the microbial rhodopsin family that directly translate absorbed light into ion fluxes along electrochemical gradients across cellular membranes by opening a conductive pore^1–3^. ChRs are composed of seven transmembrane helices and an embedded retinal cofactor linked to a conserved lysine in helix 7 via a Schiff base (retinal Schiff base, RSB). Upon photon absorption, the RSB isomerizes from all-*trans* to 13-*cis*, which induces structural changes, collectively described as spectroscopically distinguishable intermediates in a photocycle^4^.

In response to extended light pulses, the photocurrents of most known ChRs decline from an initial peak current to a lower, stationary level, a phenomenon known as desensitization (also termed inactivation)^2,4–6^. The degree and kinetics of desensitization differ among ChRs and depend on pH, membrane voltage as well as light intensity and color, with typically ≤70% amplitude reduction^1,2,7^. Photocurrent decrease via desensitization has been explained by accumulation of late non-conducting photocycle intermediates and by an alternative photocycle exhibiting low cation conductance^7–10^.

During the past fourteen years, cation-conducting ChRs (CCRs) were widely employed to depolarize genetically targeted neurons or neuronal networks using light to trigger action-potential firing^11–15^. Originally, light-driven microbial ion pumps were utilized to suppress neuronal activity by hyperpolarization^16,17^. Since ion pumps always transport one ion per absorbed photon, efficient neuronal silencing required high ion pump expression levels and continuous, intense illumination. This disadvantage was overcome by converting CCRs into anion-conducting ChRs (ACRs)^18–21^. Such engineered ACRs (eACRs)^22^ and later-discovered natural ACRs (nACRs)^23–26^ silence neuronal activity by light-induced shunting-inhibition, similar to endogenous GABA- or glycine-activated chloride channels^22,27–30^.

Here, we report a new family of phylogenetically distinct ChRs metagenomically identified from marine microorganisms. These ChRs conduct anions but exhibit unique desensitization in continuous light and were therefore designated MerMAIDs (<u>Me</u>tagenomically discovered, <u>M</u>arine, <u>A</u>nion-conducting and Intensely <u>D</u>esensitizing ChRs). Seven MerMAIDs were characterized biophysically via electrophysiological recordings, and we elucidated the molecular mechanism of the first accessible MerMAIDs using spectroscopic analyses and molecular dynamic (MD) calculations. We also explore the optogenetic inhibitory potential in neurons.

## Results

Seven putative ChRs constituting a new distinct phylogenetic branch in the ChR superfamily were identified in contigs assembled from the *Tara* Oceans metagenomes (MerMAIDs in Fig. 1a). However, the shortness of the assemblies (<10 kb) precluded taxonomic classification of the contigs. These MerMAIDs appeared to be globally distributed in the oceans, most abundant at stations near the equatorial Pacific and South Atlantic Oceans (Fig. 1b). The MerMAIDs were primarily constrained to the photic zone (depth, 0–200 m), as previously reported for other rhodopsins^31^ (Fig. S1).

**Fig. 1.**
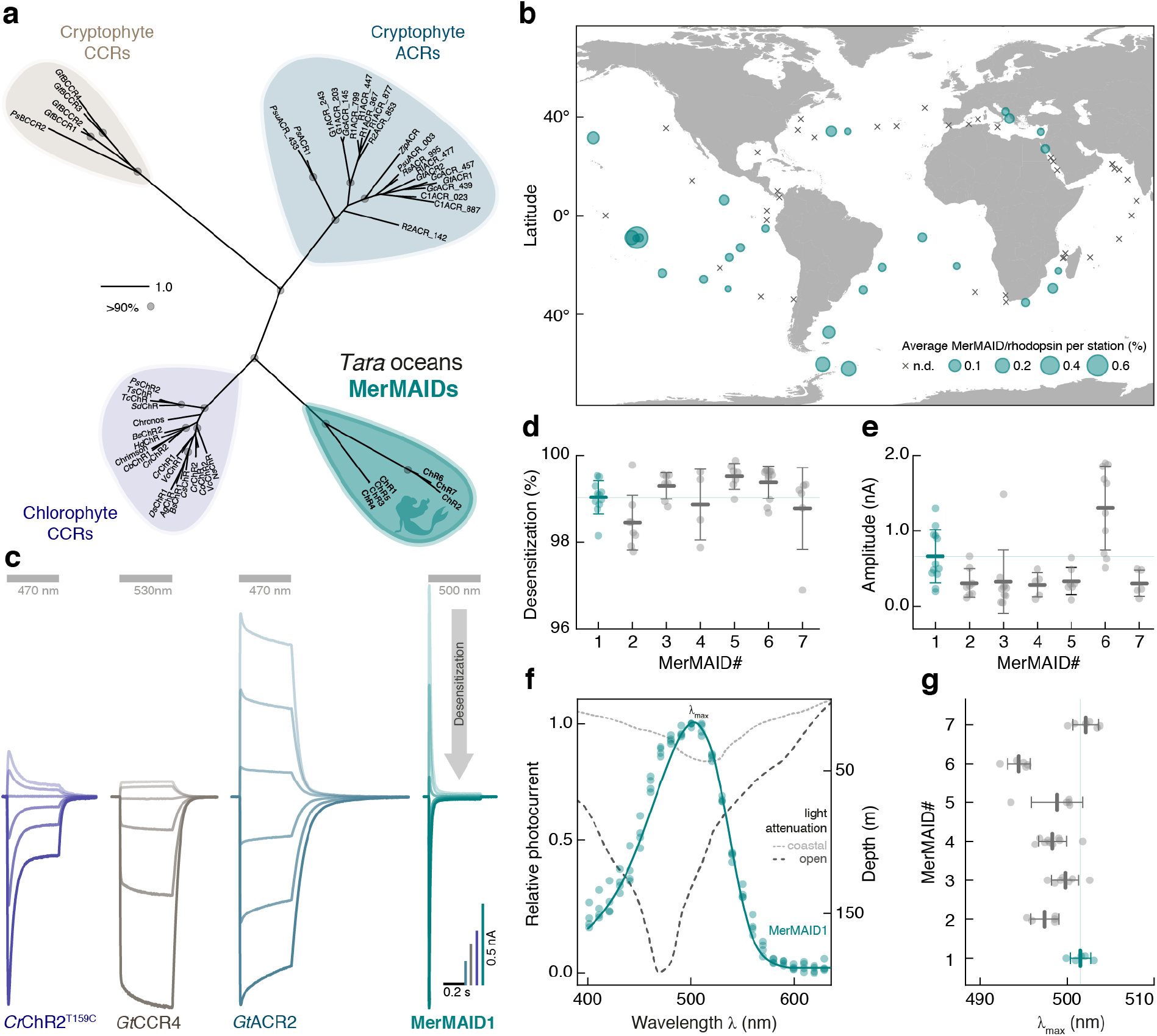
Discovery and electrophysiological features of MerMAIDs. **a**, Unrooted phylogenetic tree of the channelrhodopsin superfamily, with grey circles representing bootstrap values >90%. Scale bar indicates the average number of amino acid substitutions per site. CCR, cation-conducting channelrhodopsin; ACR, anion-conducting channelrhodopsin. **b**, Distribution and relative abundance of MerMAIDs in samples from the *Tara* Oceans project. Area of each circle indicate the estimated average abundance of MerMAID-like rhodopsins at different *Tara* Oceans stations. Stations were MerMAIDs were not detected (n.d.) are indicated by crosses. **c**, Photocurrent traces of representative members of previously identified ChR families and MerMAIDs, recorded from −60 to +40 mV in steps of 20 or 15 mV (*Gt*CCR4). Grey bars indicate light application at denoted wavelengths. **d**, **e**, Desensitization (**d**) and peak current amplitudes (**e**) of all MerMAIDs at −60 mV during continuous illumination with 500 nm light. **f**, Normalized action spectrum of MerMAID1. Single measurements are shown as dots (n=4), and solid line represents fitted data. Dashed lines indicate light penetration depth in coastal and open seawater (adopted from ref.^86^). λ_max_, maximum response wavelength; **g**, λ_max_ for all MerMAIDs. Mean values (thick lines) ± standard deviation (whiskers) are shown, and single-measurement data points are represented as dots.

Phylogenetically, the MerMAIDs appear more closely related to chlorophyte CCRs than cryptophyte ACRs (Fig. 1a). However, sequence comparisons indicated that MerMAIDs might be anion-conducting due to the lack of typical glutamate residues found in chlorophyte CCRs, though also missing in chryptophyte CCRs (Fig. S2). To examine their function, we expressed MerMAIDs in human embryonic kidney (HEK) cells and performed whole-cell voltage-clamp experiments at 1-day post transfection.

When excited with 500-nm light, MerMAID-expressing cells exhibited large photocurrents but in contrast to all previously analyzed ChRs (Fig. 1c), MerMAIDs reveal almost complete desensitization with continuous, bright light exposure (Figs. 1c,d and S3a,b). Maximum peak photocurrent amplitudes reached up to 2 nA (MerMAID-6, Figs. 1e and S3c), but the current did not saturate even at 10 mW/mm^2^ (Fig. S3d). Transient photocurrent action spectra were recorded to determine the wavelength sensitivity of the MerMAIDs. All variants tested exhibited typical rhodopsin spectra, with maximal sensitivity close to 500 nm (Figs. 1f,g and S3e), as expected for marine organisms, given that blue light penetration is strongest within the photic zone in seawater (Fig. 1f).

Next, we tested the ion selectivity of the MerMAIDs. Because we suspected anion selectivity, we depleted the extracellular Cl^−^ from 150 mM to 10 mM while maintaining the intracellular Cl^−^ at 120 mM (Fig. 2a). This increased the inward current and induced an upshift of the reversal potential (Fig. 2a,b), consistent with a Cl^−^ efflux. A similar shift close to the calculated Cl^−^-Nernst potential was obtained for all MerMAIDs (Figs. 2c and S4a), as well as for the small stationary photocurrents of MerMAID1 (Figs. 2a,c and S4a). These data justified the classification of MerMAIDs as ACRs.

**Fig. 2.**
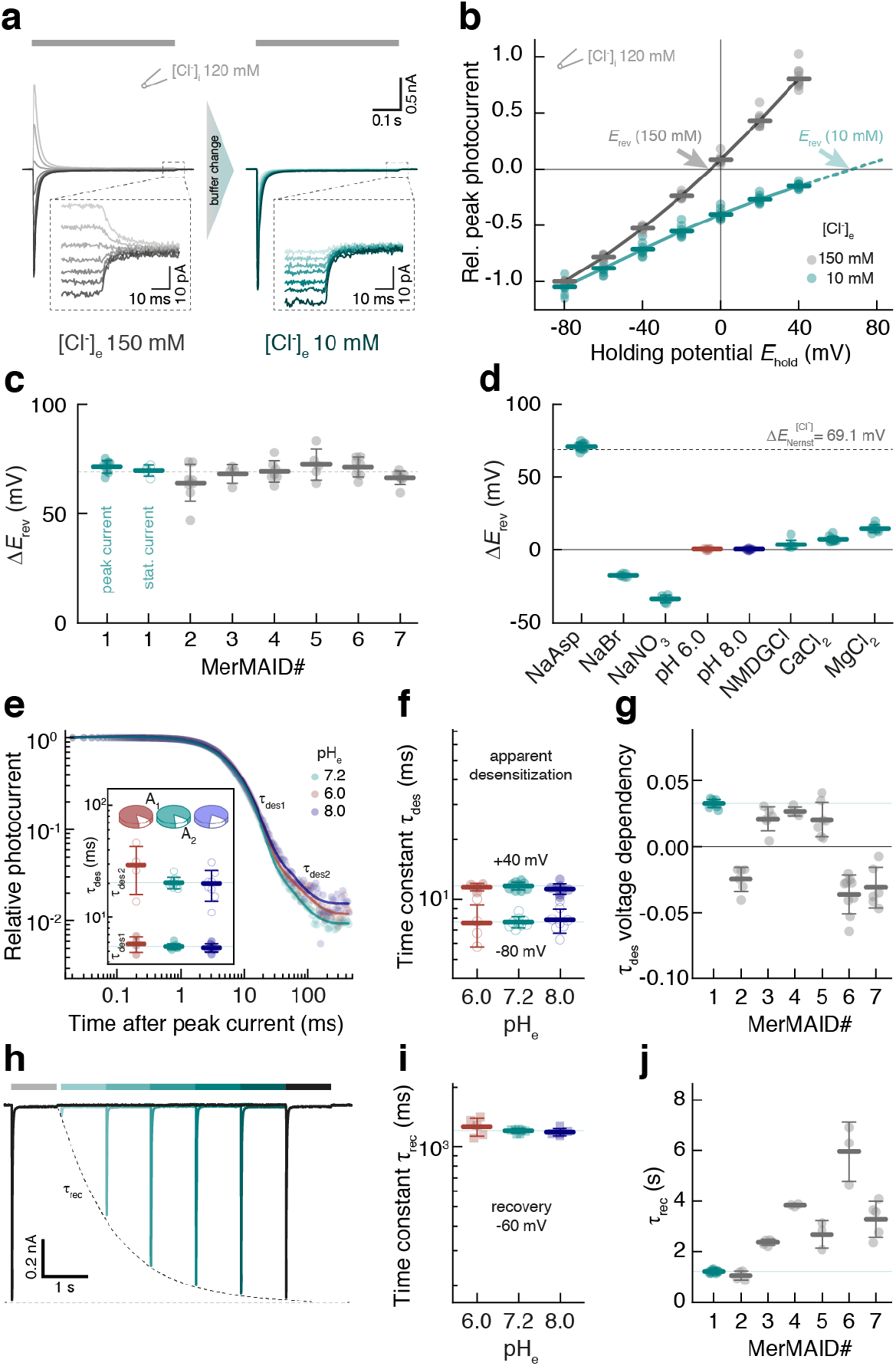
Ion selectivity and kinetic properties of MerMAIDs. **a**, Representative photocurrent traces of MerMAID1 elicited with 500 nm light (grey bar) at different holding potentials (−80 to +40 mV, in 20-mV steps, from bottom to top) before (left, grey) and after extracellular chloride reduction (right, cyan), as indicated. Insets show enlarged views of the remaining stationary photocurrent. **b**, Current-voltage relationship of the MerMAID1 peak photocurrent at 150 mM (grey) and 10 mM (cyan) extracellular chloride ([Cl^−^]_e_). Arrows indicate reversal potentials (*E*_rev_). **c**, Reversal potential shifts (Δ*E*_rev_) upon reduction of [Cl^−^]_e_ for peak currents of all MerMAIDs as well as the stationary current of MerMAID1. **d**, Δ*E*_rev_ values of MerMAID1 upon exchange of external buffer. Δ*E*_*r*ev_ of the theoretical Nernst potential for Cl^−^ is indicated as a dashed line (**c**, **d**). **e**, Extracellular pH (pH_e_) dependence of biphasic MerMAID1 desensitization kinetic. Inset shows the time constants and their relative amplitudes to total decay. **f**, pH_e_ dependency of the apparent desensitization time constant (τ_des_) at −80 and +40 mV. **g**, Voltage dependency of τ_des_ for all MerMAIDs in ms/mV. **h**, Double-light pulse experiment at −60 mV and pH_e_ 7.2 to determine the peak current recovery time constant (τ_rec_). **i**, pH_e_ dependency of MerMAID1 at −60 mV. **j**, Recovery time constants of all MerMAID variants. Mean values (thick lines) ± standard deviation (whiskers) are shown, and single-measurement data points are represented as dots.

To evaluate the conductance of other anions, we performed ion substitution experiments using MerMAID1 as a model. Replacement of Cl^−^ with Br^−^ or NO_3_^−^ resulted in negative reversal potential shifts (Figs. 2d and S4b), thus revealing nonselective anion conductivity, as previously reported for other ACRs^21,23^. In contrast, substitution of Na^+^ with NMDG^+^, Ca^2+^, or Mg^2+^ had only a slight effect on reversal potentials (Fig. 2d and S4b), thereby excluding a substantial contribution by cations as charge carriers.

Rhodopsin function often commonly involves de- and reprotonation of internal amino acids, and pH changes can significantly affect photocurrent amplitude and kinetics^4,9,18,19^. We therefore investigated the effect of extra- and intracellular pH (pH_e_ and pH_i_) changes on MerMAID1. Variation of pH_e_ between 6.0 and 8.0 slightly altered the photocurrent amplitude (Fig. S4c) but not the reversal potential (Figs. 2d and S4d), thus excluding proton transport. Neither pH_i_ nor pH_e_ affected the desensitization time constant, τ_des_ (Figs. 2e,f and S4e-g). However, τ_des_ exhibit a moderate voltage dependence (Figs. 2f,g and S4e,f). For MerMAID1,3-5, desensitization slowed down with increasing holding potential whereas τ_des_ was accelerated for MerMAIDs 2, 6, and 7 (Fig. 2g). These groups correlated well with the two phylogenetic branches within the MerMAID family (Fig. 1a), although the underlying molecular determinants of this difference remain unknown.

To assess the photocycle turnover time (recovery kinetic time constant, τ_rec_), we performed double-pulse measurements at −60 mV (Fig. 2h) at different pH_e_ values. The peak current recovered with τ_rec_=1.21 ± 0.03 s for MerMAID1, and was unaffected by pH_e_ or pH_i_ changes (Figs. 2i,j and S4h,i). Between the different MerMAIDs, τ_rec_ varied between 1.1 ± 0.2 s (MerMAID2) and 6 ± 1 s (MerMAID6, Figs. 2j, and S4i).

To elucidate the desensitization mechanism, recombinant MerMAID1 was purified from *Pichia pastoris* and analyzed by UV/vis and vibrational spectroscopy. Steady-state UV/vis absorption spectra of dark-adapted MerMAID1 exhibited a prominent peak at 502 nm, consistent with the photocurrent action spectra (Fig. 3a). Upon continuous illumination with green light, the 502-nm dark-state absorption peak decreased, while a fine-structured, blue-shifted intermediate with sub-maxima at 346, 364, and 384 nm accumulated in parallel (Fig. 3a,d). Similarly, alkalization converted dark-adapted MerMAID1 into a more red-shifted, fine-structured UV-absorbing species, consistent with a deprotonated 13-*cis* isomer in the M-state and deprotonated all-*trans* RSB dark state^32^ that occurs with a pK value of approximately 9.8 (Figs. 3b,d and S5d).

**Fig. 3.**
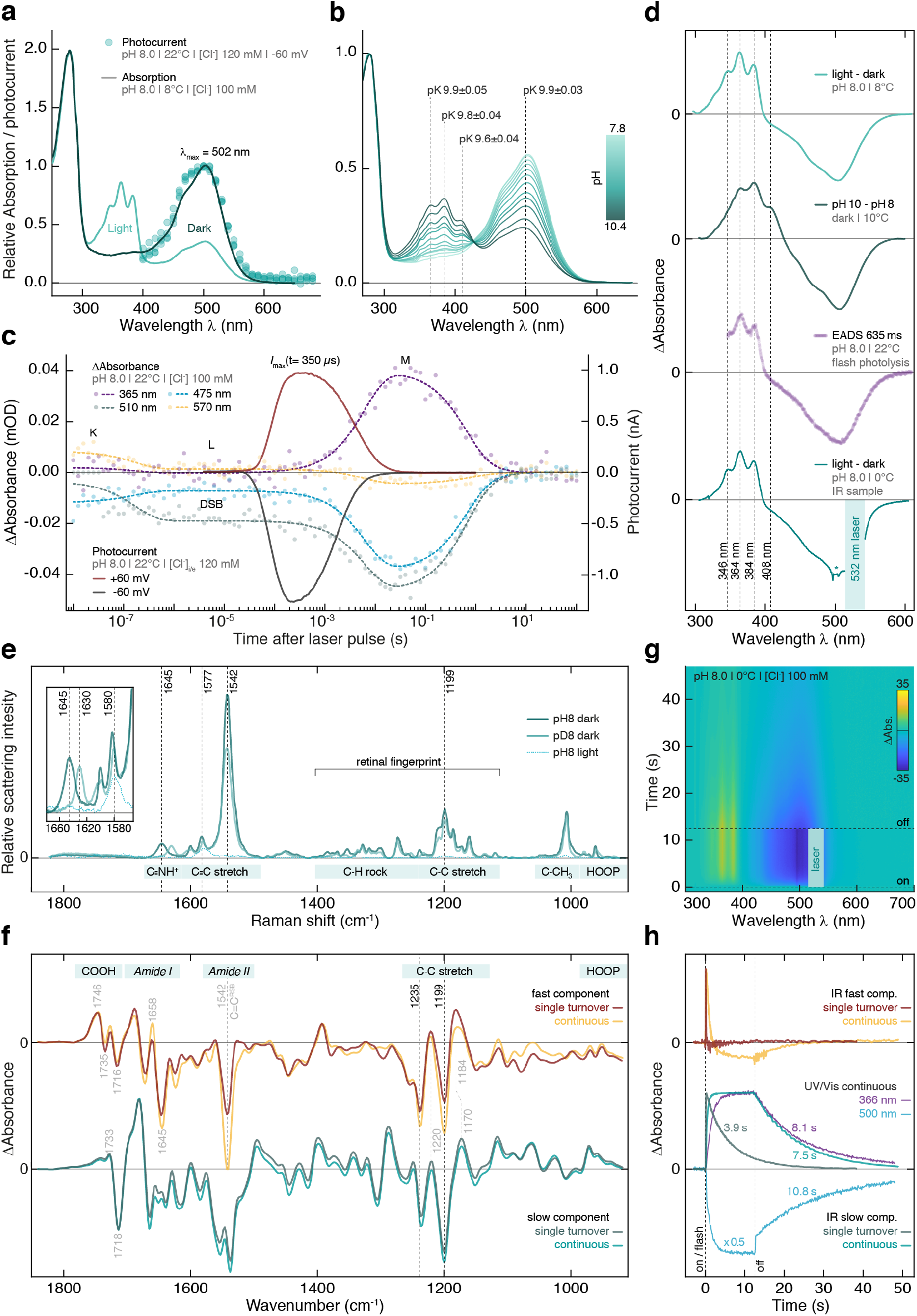
Spectroscopic characterization of purified MerMAID1. **a**, Normalized UV/vis absorption spectra of dark-adapted and illuminated MerMAID1. Filled circles indicate single-measurement action spectra recordings, as shown in Figure 1f. **b**, Normalized UV/vis absorption spectra of MerMAID1 at different pH values, titrated from pH 7.8 to 10.4. The pK values for specific wavelengths are indicated. **c**, Transient absorption changes and electrophysiological recordings obtained with single-turnover laser pulse excitation. **d**, Fine-structured difference absorption spectra obtained from different experiments. (From top to bottom) light minus dark difference spectra obtained from data shown in panel a, pH-difference spectra calculated from panel b data, evolution-associated difference spectra (EADS) resulting from a global fit of the transient absorption spectra and (bottom) light-minus-dark difference spectrum measured using the FTIR sample shown in (**g**). Due to strong laser scattering, a portion of the spectral data is excluded for the FTIR sample, and residual scattering is marked with an asterisk. **e**, Resonance Raman spectra of dark-adapted MerMAID1 at pH/D 8 (recorded at 488 nm) as well as cryo-trapped and illuminated protein sample at pH 8 (recorded at 413 nm). Inset: zoomed C=NH^+^ stretching region. **f**, Kinetically decomposed FTIR light-minus-dark absorption of MerMAID1, recorded with single turnover and continuous illumination at 0°C. Bands marked in grey are discussed in the Supplementary Material **g**, Contour plot of transient absorption changes of the sample used in (**f**) illuminated with a 532 nm continuous laser. **h**, Decay kinetics of the fast and slow FTIR components obtained under single-turnover and continuous illumination conditions, respectively. Kinetics at 366 and 500 nm obtained from the UV/vis spectroscopic measurements shown in (**g**) are shown for comparison.

Single-turnover voltage-clamp experiments showed a maximum channel conductance 350 µs after ns-pulse laser excitation. Channel closing was biphasic, with a dominant fast component and an apparent closing time constant (τ_off_) of 2.7 ± 0.1 ms (Figs. 3c and S5a). Transient UV/vis absorption spectra (Figs. 3c and S5b,c) revealed an early decaying (173 ns) K-like photoproduct observed only briefly on our time scale. The evolution-associated difference spectrum (EADS) of the subsequent L-intermediate is slightly blue shifted and to some extent broadened compared to the dark-state spectrum (Fig. S5c), indicating closer proximity of the primary counterion to the RSBH^+^ immediately prior to RSB deprotonation^33^. Within 6 ms, the L-state converted to the M-state, with concomitant deprotonation of the RSB, as indicated by the large blue shift coinciding with channel closure (Figs. 3c and S5a,b). The transient M-state EADS was similarly fine-structured as observed for continuous photo activation (Fig. 3d), indicating accumulation of the M-state during sustained light exposure.

To assess potential retinal chromophore isoforms of MerMAID1, we performed resonance Raman (RR) spectroscopy at 80 K. Excitation of dark-adapted MerMAID1 at 488 or 514 nm produced identical RR spectra (Fig. S5e), indicating structural homogeneity of the chromophore. The vibrational band pattern in the retinal fingerprint region (1100–1400 cm^−1^) was characteristic of an all-*trans* RSB^34,35^ (Fig. 3e). Upon proton/deuterium exchange, the C=N stretching mode downshifted from 1645 to 1630 cm^−1^ (inset Fig. 3e), indicative of a weakly hydrogen-bonded^36^ protonated RSB^34,35^. RR spectra of photoactivated MerMAID1 cryo-trapped in the M-state and excited at 413 nm exhibited a prominent band at 1577 cm^−1^ attributable to a 13-*cis* configuration of the chromophore with deprotonated RSB^37^. RR spectra of dark-adapted MerMAID1 probed with 488 or 413 nm at pH 10 were similar to spectra of dark- and light-adapted MerMAID1 at pH 8 (Fig. S5e). Notably, strong bands in the C=C stretching region at ca. 1540 and 1577 cm^−1^ indicated a mixture of two chromophore isomers, corresponding to the protonated and deprotonated all-*trans* RSB of the dark state at high pH. Thus, RR spectra confirmed that MerMAID1 undergoes all-*trans* to 13-*cis* retinal isomerization and deprotonation of the RSB during illumination and may deprotonate at high pH in the dark.

Time-resolved Fourier-transform infrared (FTIR) spectra were collected at pH 8.0 and 0 °C to examine the light-driven molecular processes of MerMAID1 under single-turnover conditions and continuous illumination (Fig. 3f,h), with parallel UV/vis observation of M-state formation (Fig. 3g, h). Kinetic decomposition of light-dark FTIR difference spectra revealed highly similar fast and slow spectral components for both single-turnover and continuous illumination, respectively (Fig. 3f). At both conditions, the slow FTIR component relaxed to the dark state mono-exponentially, within seconds (Fig. 3h) and was assigned to the late M-state that accumulated with continuous illumination (Fig. 3g,h) without formation of other photoproducts. This assignment was supported by data for the retinal fingerprint region that—similar to the RR data—indicated all-*trans* to 13-*cis* retinal isomerization as the only photoreaction based on the negative bands at 1235(−) and 1199(−) cm^−1^. The fast FTIR spectral components resembled the short-lived conducting L-state preceding the late M-intermediate, as inferred from the comparable decay kinetics (see Supplemental Discussion).

Site-directed mutagenesis guided by MD simulations and probed by electrophysiological recordings were conducted to further examine the molecular mechanism for the intense desensitization of the MerMAIDs. For MD simulations, a MerMAID1 homology model was constructed based on the iC++ crystal structure^38^ and embedded in a phospholipid bilayer (Figs. 4a and S6a). The D210, E44, W80, and Y48 side chains located near the protonated Schiff base (Fig. 4a,c) maintained their relative positions during a 100-ns MD simulation. The orientation and distances of these residues changed only slightly with inflowing water (Fig. S6d-f). Possible ion translocation pathways were calculated using MOLEonline^39^. Figure 4a,b shows the most likely ion pathway based on surface charge considerations (Fig. S6b,c). Extracellularly, MerMAID1 is accessible via a narrowing tunnel that is disrupted by W80, D210, and the RSB (Fig. 4a,b). Intracellularly, another ion pathway is formed leading from the protein surface almost to the Schiff base, disconnected only by a short hydrophobic barrier. In our model, the carboxylic residues of the active site complex (E44 and D210) were deprotonated based on pK_a_ calculations (Fig. 4c, pK_a_ <5.5). D210, which acts as closest counterion (2.6 Å) to the Schiff base nitrogen, primarily stabilizes the RSB proton (Fig. 4c). The carboxyl group of D210 also interacts with S79, Y48, and E44 via two water molecules, whereas E44 hydrogen bonds directly to Y48 and is linked to the backbone oxygen of D210 via another water molecule (Fig. 4c). When the counterion D210 is neutralized (D210N), photocurrents are drastically reduced (Fig. 4d,e), λ_max_ was 16 nm red-shifted (Fig. 4f), and the recovery kinetics decelerated markedly (Fig. 4h). Elimination of the more distant E44 via an E44Q mutation caused only a 3 nm bathochromic action-spectrum shift (Fig. 4f), increased the photocurrent amplitudes (Fig. 4f), decelerated desensitization by a factor of 10 (Fig. 4d,g) and slightly reduced the extent of desensitization (Fig. 4i). Replacement of both acidic residues (E44Q-D210N) only halved photocurrent amplitudes (Fig. 4e) and shifted λ_max_ to 513 ± 1 nm (Fig. 4f), suggesting rearrangement of the hydrogen bond network around the RSB. Desensitization remained strong (Fig. 4i), but the kinetics slowed, similar to the E44Q mutation alone (Fig. 4g).

**Fig. 4.**
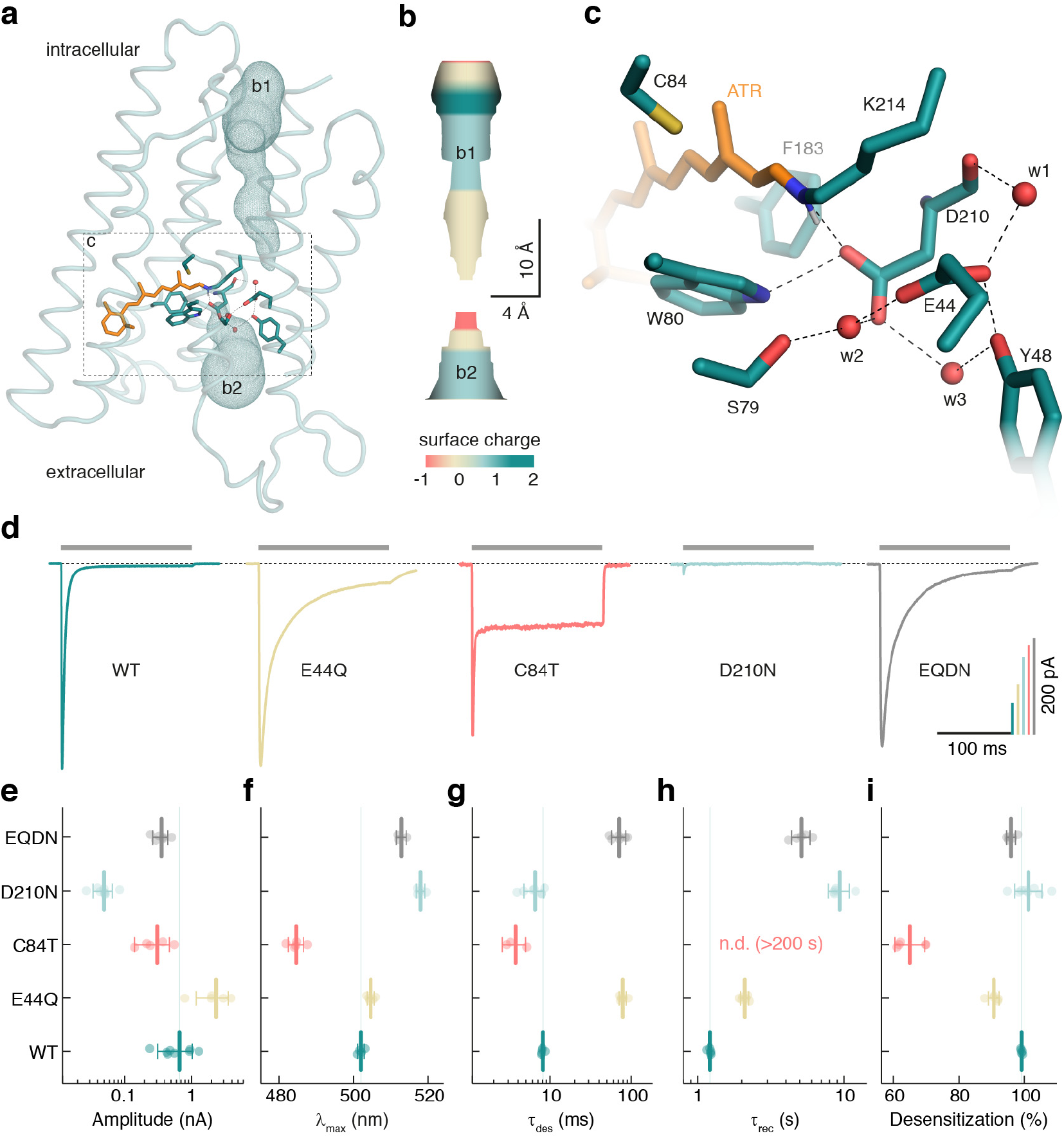
MD simulations and mutational analysis of MerMAID1. **a**, Overview of the MD simulation homology model of MerMAID1 in the dark. The predicted ion permeation pathway is shown as mesh (b1, b2), and ribbons represent the protein backbone. **b**, Electrostatic surface potential of the predicted chloride permeation pathway. **c**, Detailed view of the active-site residues, with amino acids shown as cyan sticks and the all-*trans* retinal (ATR) in orange. Red spheres denote water molecules that remained stable during MD simulation. **d**, Representative photocurrent traces of wild-type (WT) MerMAID1 and selected MerMAID1 mutants recorded at −60 mV. Photocurrent amplitudes (**e**), λ_max_ (**f**), apparent τ_des_ of the peak current (**g**), recovery time constant, τ_rec_ (**h**), and extent of desensitization (**i**) of WT MerMAID1 and indicated mutants. Mean values (thick lines) ± standard deviation (whiskers) are shown, and single-measurement data points are represented as dots.

Neutralization of E44 increased the stationary current only slightly (Fig. 4i), whereas we identified C84 (the *Cr*ChR2 C128 homolog) as a crucial determinant of the inactivation process. The C84T mutant exhibited a decreased peak current amplitude but markedly increased stationary photocurrent (Fig. 4d,e), resulting in only 65 ± 5 % desensitization (Fig. 4d,i) and minimally altered desensitization kinetics (Fig. 4d,g). In contrast, we observed no peak current recovery within a time period of 200 s. As suggested by our model structure and the pronounced 17 nm blue-shifted λ_max_ (Fig. 4f), C84 is located near the retinal polyene chain and the C13 methyl group (Fig. 4a,c). In MerMAIDs, this cysteine cannot serve as a link between helices 3 and 4 as discussed for bacteriorhodopsin^40,41^ and CCRs^42,43^ due to the absence of a hydrogen-bonding partner in helix 4 (Fig. S2). C84 might stabilize the deprotonated RSB within the anion permeation pathway instead, thereby disrupting further ion conduction via charge repulsion of the deprotonated RSB during the photocycle.

Finally, we evaluated the utility of MerMAIDs as optogenetic tools for inhibiting neuronal activity. As MerMAID6 exhibited the highest photocurrent in HEK cells (Fig. 1e), we generated a Citrine-labeled MerMAID6 variant and co-expressed it with mCerulean as a volume marker. MerMAID6-Citrine expression was readily detected in CA1 pyramidal neurons of hippocampal slice cultures 4 to 5 days after single-cell electroporation. We observed membrane-localized MerMAID6 expression, with some fraction of the protein displaying a speckled cellular distribution (Fig. 5a). However, illumination triggered high transmembrane photocurrents with biophysical properties similar to those observed in HEK cells (Fig. S8). The large, transient photocurrents observed in neurons led us to hypothesize that MerMAID6 could be used to block single action potentials (APs) with high temporal precision and without affecting subsequent APs in the presence of light. We first injected a depolarizing current ramp into the soma to precisely determine the rheobase for AP firing in the dark. A 10 ms light pulse synchronized with the first AP that occurred during darkness eliminated generation of this AP (Fig. 5b). We then applied a 500 ms light pulse synchronized to the time of onset of the first AP lasting throughout the remainder of the current ramp (Fig. 5c) or a depolarizing current step (Fig. 5d), MerMAID6 suppressed generation of the first AP, without affecting the following ones due to rapid accumulation of the desensitized and non-conducting state during extended illumination. Similarly, selective inhibition of a single AP was achieved with MerMAID1 (Fig. S7), demonstrating that photoactivated MerMAID1 and 6 provide efficient and temporally precise inhibition of neuronal activity.

**Fig. 5.**
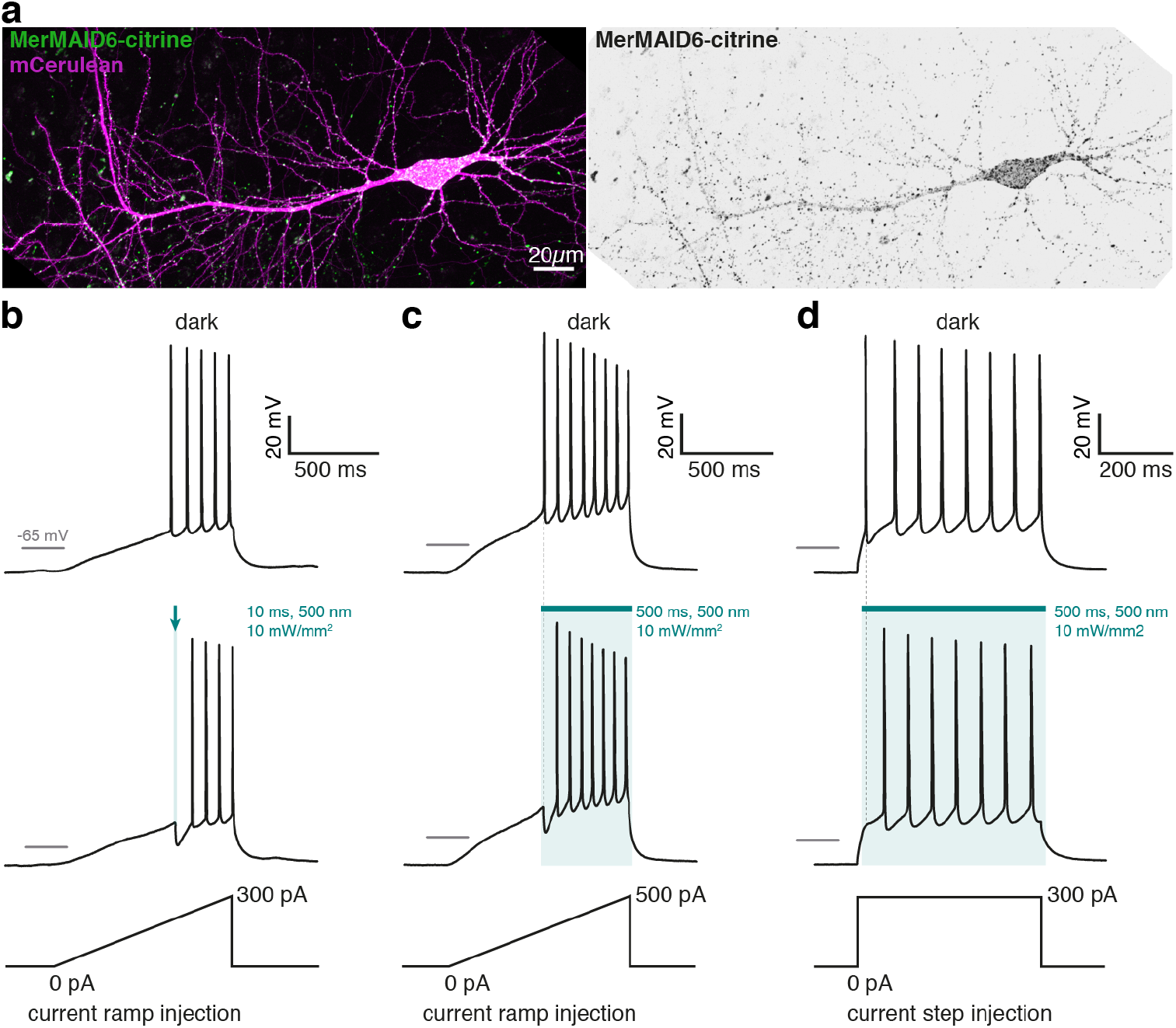
Neuronal application of MerMAID6 as optogenetic silencer. **a**, CA1 pyramidal neuron expressing MerMAID6-Citrine (green) 5 days after electroporation (stitched maximum intensity projections of two-photon images). mCerulean (magenta) was co-electroporated to visualize neuronal morphology (left). Fluorescence intensity shown as inverted gray values (right). **b** and **c**, Voltage traces in response to depolarizing current ramps injected into MerMAID6-expressing CA1 pyramidal cells. Illumination with green light (500 nm, 10mW/mm^2^) for a brief (10 ms, **b**) or longer (500 ms, **c**) time period blocked single spikes. Light onset preceded action potential onset (measured in the dark condition) by 5 ms. **d**, Same as (**c**) but a depolarizing current step of 300 pA was injected into the neuron instead of a current ramp.

## Discussion

We extensively characterized the biophysical properties of the MerMAIDs, a new family of ChRs identified from metagenomic data. All MerMAIDs share similar activity maxima optimal for sensing light in moderate-depth seawater. Similar to other recently discovered natural ACRs^23^, MerMAIDs selectively conduct anions. Distinct from all other ChRs, MerMAIDs exhibit almost complete desensitization during exposure to continuous bright light. However, the environmental advantage of near-complete desensitization compared with non-inactivating ACRs is unclear.

After photon absorption, the MerMAID1 chromophore isomerizes from all-*trans* to 13-*cis*, as demonstrated by RR and FTIR spectroscopy. We hypothesize that the RSBH^+^ dipole changes orientation and distance with respect to the nearby D210, as evidenced by formation of the L-intermediate, analogous to bacteriorhodopsin^33^. The K→L transition and formation of the open state is accompanied by minimal protein backbone changes that induce changes in hydrogen bonding and side-chain pK values, presumably involving the active-site residues D210 and E44. Maximum channel conductivity, reached within 350 µs, is UV/vis spectroscopically almost silent. Channel closing proceeds concurrently with RSB deprotonation, leading to M-intermediate formation, similar to cryptophyte ACRs^24,44^ but in contrast to chlorophyte CCRs, in which M-state formation precedes channel opening^45,46^. These observations suggest that the positively charged protonated RSB is part of the chloride conducting pathway, consistent with the calculated ion permeation pathway along the counter-ion complex, similar to crystal structures of *Guillardia theta* ACR1 (*Gt*ACR1)^38,47^. Chloride flux in MerMAID1 is interrupted by charge repulsion following deprotonation of the Schiff base. In the final photocycle step, MerMAID1 structurally rearranges, the RSB reprotonates, and the initial dark state is fully repopulated within seconds.

The observation that photocurrent kinetics were not affected by intra- or extracellular pH changes suggests that the RSB proton remains within the central active-site complex during the photocycle, as recently reported for heliorhodopsins^31^. D210 is the primary counterion of the RSB in MerMAID1, but both carboxylic residues (E44 and D210) participate in de- and reprotonation of the MerMAID1 chromophore, as neutralization of either one or both residues affects formation of the conductive or desensitized state. However, retention of function of the E44Q-D210N double mutant suggests the possibility of alternative proton acceptor and donor sites.

The unique desensitization of the MerMAIDs can be explained by the accumulation of the blue-shifted M-intermediate during constant photoactivation. Because the non-conducting M-intermediate is formed within milliseconds and decays only within hundreds of milliseconds, the current declines to 1 % in continuous light. This mechanism is consistent with an M-state that cannot be photochemically converted back to the dark state (Fig. S5g,h). As discussed in previous reports, decline of *Cr*ChR2 photocurrents upon continuous illumination is due to both, the accumulation of late non-conducting photocycle intermediates and population of a parallel (*syn-*) photocycle with an only weakly conducting open state^7–9^. The accumulation of a late photocycle intermediate is the dominant mechanism in MerMAID1, as demonstrated by FTIR spectroscopy; no parallel photocycle is needed to explain the strong inactivation.

We found that replacing C84 in MerMAID1 decreases the peak photocurrent but increases stationary photocurrents, thus reducing the extent of desensitization, possibly due to either a prolonged L-state or a shortened recovery from the M- to the initial dark state. C84 could have dual functions in wild-type MerMAID1: i) stabilizing the deprotonated RSB to retain the repulsing charge in the ion pathway and block the channel; ii) suppression of C=N *anti* to *syn* isomerization and population of parallel *syn*-photocycles as discussed for *Cr*ChR2^10^.

Another unusual feature of MerMAID1 are the fine-structured absorption spectra of both the deprotonated all-*trans* RSB in the dark at alkaline pH and the 13-*cis* retinal of the M-state. Such unusual spectra have been reported for other microbial rhodopsins after retro-retinal formation upon reduction with borohydride^48^ or hydrolysis of the RSB^49^. In both cases, the UV fine structure results from immobility of the deprotonated chromophore, which is typically more pronounced at deep temperatures^50,51^. Alkalization-induced fine-structured spectra were reported for eACRs^38^ and wild-type and mutant nACRs^44^ and suggested to result from RSB hydrolysis^38^ or protein denaturation^52^. However, in *Gt*ACR1, the covalent bond between the retinal and the Schiff base– forming lysine is not broken at high pH. Instead, the RSB deprotonates and adopts an M-like configuration^53^. RR spectra of MerMAID1 at pH 10 (Fig. S5e) suggested that the retinal is similarly deprotonated and adopts a rigid configuration in all-*trans* instead of 13-*cis*, as in the M-state.

MerMAID1 and 6 effectively inhibited neuronal activity with high temporal precision. Due to the unique desensitization of the MerMAIDs under continuous illumination, single APs can be blocked at the onset of illumination without affecting subsequent neuronal spiking. Hence, MerMAIDs could serve as transient optogenetic silencers to inhibit individual APs with high precision in combination with subsequent imaging of spectrally overlapping reporters of neuronal activity. MerMAIDs would thus facilitate continuous monitoring of neuronal activity subsequent to short-duration inhibition at the same wavelength.

The identification of the entirely new ChR family fortifies the value of metagenomic data for the discovery of novel photoreceptor proteins potentially applicable as optogenetic tools. The initial in-depth characterization of MerMAIDs will foster the generation of ChRs with novel biophysical properties and lead to deeper understanding of the working principles of rhodopsins.

## Methods

### Identification of novel channelrhodopsins and metagenomics data analysis

Novel channelrhodopsin variants were identified using full-length *Cr*ChR1 and *Cr*ChR2 amino acid sequences (GenBank: AAL08946.1, and NCBI reference sequence: XP_001701725.1, respectively) as queries for tblastn 2.6.0 analysis^54^ against a database of contigs assembled from the *Tara* Oceans metagenomic datasets of bacterial^55^, viral^56^, and girus^57^ samples. The assemblies were generated as described elsewhere^58^.

MerMAID abundance in the marine environment was estimated using the Ocean Gene Atlas^59^ after mining the Ocean Microbial Reference Gene Catalog^55^. A collection of representative microbial rhodopsin protein sequences from distinct subfamilies containing the MerMAIDs was aligned using the MAFFT online server (ver. 7)^60^. The alignment was used to generate a Hidden Markov Model (HMM) using hmmbuild from the HMMER 3.1b2 suite^61^. The HMM served as the query in the Ocean Gene Atlas^59^ or an HMMER-based search with default parameters against the Ocean Microbial Reference Gene Catalog. The Ocean Gene Atlas results for abundances and homologs were stored locally for further analysis.

Protein homologs from the Ocean Gene Atlas and MerMAIDs were pooled and aligned using the MAFFT web server. MAFFT multiple sequence alignment was used to identify those homologs phylogenetically closer to the MerMAIDs and were tagged as MerMAID-like. Ocean Gene Atlas abundance data were parsed using a custom R script to calculate the ratio of ACR-like proteins to total rhodopsins in each *Tara* Oceans sample. The MerMAID-like/total rhodopsin ratio was coupled with environmental metadata from the *Tara* Oceans samples to generate depth profiles and distribution maps using the R packages maps^62^, ggplot2^63^, and ggalt^64^.

The phylogenetic tree was generated using phylogeny.fr^65^ and the sequence alignment using Clustal X^66^. The sequence alignment was visualized using the ENDscript 2 web server^67^, and the alignment was cropped to include the transmembrane regions of selected ChRs.

### Molecular biology and protein purification

Human/mouse codon-optimized sequences encoding MerMAIDs were synthesized (GenScript, Piscataway, NJ) and cloned into the p-mCherry-C1 vector using *Nhe*I and *Age*I restriction sites (FastDigest, Thermo Fisher Scientific, Waltham, MA) for electrophysiologic recordings in HEK293 cells. Due to incomplete metagenomic data, a methionine was added as start codon for MerMAID1 and MerMAID4. Site-directed mutagenesis was performed using *Pfu* polymerase (Agilent Technologies, Santa Clara, CA). MerMAID1 and MerMAID6 cDNAs were subcloned into neuron-specific expression vectors (pAAV backbone, human synapsin promoter) in frame with Citrine cDNA using Gibson assembly^68^. For expression in *Pichia pastoris*, the MerMAID1 gene was subcloned with a C-terminal TEV protease restriction site and a 6× His-Tag into the pPiCZ vector (Invitrogen, Carlsbad, CA). Zeocin™-resistant positive clones were selected from electroporation-transformed yeast cells. Expression of MerMAID1 in precultured cells was induced with 2.5% methanol in presence of 5 µM all-*trans* retinal for 24 h. Cells were harvested by centrifugation and resuspended in breaking buffer (50 mM NaPO_4_, 1 mM EDTA, 1 mM PMSF, 5% glycerol [pH 7.4]) and disrupted by high pressure using a French press (G. Heinemann Ultraschall und Labortechnik, Schwäbisch Gmünd, Germany). The membrane fraction was collected, homogenized, and solubilized overnight at 4°C in 100 mM NaCl, 20 mM Tris-HCl, 20 mM imidazole, 1 mM PMSF, 5 µM all-*trans* retinal, and 1% (*w/v*) dodecyl maltoside (DDM). Recombinant rhodopsin was purified by affinity chromatography (HisTrap™ FF Crude column, GE Healthcare Life Science, Chicago, IL) and gel filtration (HiPrep™ 26/10 desalting, GE Healthcare Life Science). Before elution, an additional washing step with buffer containing 50 mM imidazole was performed. Purified protein was concentrated and stored in 100 mM NaCl, 20 mM Tris-HCl (pH 8), and 0.05% DDM.

### Electrophysiology in HEK293 cells

HEK293 cell culture and electrophysiologic experiments were performed as described previously^22,69^. Cells were supplemented with 1 µM all-*trans* retinal, seeded at a density of 1 × 10^5^/ml on poly-D-lysine–coated coverslips, and transiently transfected using Fugene HD (Promega, Madison, WI). At 1 to 2 days post-transfection, whole-cell patch-clamp recordings were performed at 24 °C. The resistance of fire-polished patch pipettes was 1.5–2.5 MΩ, and a 140 mM NaCl agar bridge served as the reference electrode. Membrane resistance was generally ≥1 GΩ, and the access resistance was <10 MΩ. Signals were amplified (AxoPatch200B), digitized (DigiData400), and acquired using Clampex 10.4 (all from Molecular Devices, Sunnyvale, CA). Light from a Polychrome V (TILL Photonics, Planegg, Germany) with 7 nm bandwidth was channeled into an Axiovert 100 microscope (Carl Zeiss, Jena, Germany) controlled via a programmable shutter system (VS25 and VCM-D1; Vincent Associates, Rochester, NY). Light intensity was measured in the sample plane using a calibrated optometer (P9710; Gigahertz Optik, Türkenfeld, Germany) and calculated for the illuminated field of the W Plan-Apochromat 40×/1.0 DIC objective (0.066 mm^2^, Carl Zeiss). Final buffer osmolarity was set with glucose to 320 mOsm (extracellular) or 290 mOsm (intracellular), and the pH was adjusted using N-methyl-D-glucamine or citric acid. Liquid junction potentials were calculated (Clampex 10.4) and corrected. For ion selectivity measurements, extracellular buffers (supplementary table S1) were exchanged in random order by adding at least 3 ml to the measuring chamber (volume ∼0.5 ml). Fluid level was kept constant using an MPCU bath handler (Lorentz Messgerätebau, Katlenburg-Lindau, Germany). MerMAID photocurrents were induced for 500 ms and recorded between −80 and +40 mV in 20-mV steps. Low-intensity light between 390 and 680 nm was applied in 10-nm steps for 10 ms at −60 mV to generate action spectra. Equal photon irradiance at all wavelengths was achieved using a motorized neutral-density filter wheel (Newport, Irvine, CA) in the light path, controlled by custom software written in LabVIEW (National Instruments, Austin, TX). For light titration experiments, photocurrents were induced for 2 s at −60 mV, and light was attenuated using ND filters (SCHOTT, Mainz, Germany) inserted into the light path using a motorized, software-controlled filter wheel (FW212C, Thorlabs, Newton, NJ). Single-turnover experiments were performed with the above mentioned setup described elsewhere^10,70^. An Opolette HE 355 LD Nd:YAG laser/OPO system (OPOTEK, Carlsbad,CA) served as pulsed laser light source.

### UV-Vis Spectroscopy

Steady-state absorption spectra were recorded using a Cary 300 UV/vis spectrophotometer (Varian Inc., Palo Alto, USA) or UV-2600 UV/vis spectrophotometer (Shimadzu, Kyōto, Japan) at a spectral resolution of 1 nm in buffer containing 100 mM NaCl, 20 mM Tris, and 0.05% DDM (pH 8). Light-adapted absorption spectra were acquired by illuminating the sample with a 530 nm LED with a 520 ± 15 nm filter. For pH titration experiments, small volumes of 1 M NaOH were added to samples in titration buffer (100 mM NaCl, 10 mM BTP, 10 mM CAPS, 0.05% DDM [pH 7.5]). The pH was measured using pH microelectrodes (SI Analytics, Mainz, Germany). The transient absorption spectroscopic measurements shown in Figure S5b,c were performed as previously reported^24^ at 22°C using a modified LKS.60 flash-photolysis system (Applied Photophysics Ltd., Leatherhead, UK). For sample excitation, the laser pulse was tuned to 500 nm using a tunable optical parametric oscillator (MagicPrism, OPOTEK), which was pumped with the third harmonic of a Nd:YAG laser (BrilliantB, Quantel, Les Ulis, France). The laser energy was adjusted to 5 mJ/shot and pulse duration of 10 ns. A 150-W xenon lamp (Osram, München, Germany) was used to monitor changes in absorption. Transient spectra were recorded in multi-wavelength data sets at a resolution of 0.4 nm using an Andor iStar ICCD camera (DH734; Andor Technology Ltd, Belfast, Ireland). Spectra were recorded at 101 different time points between 10 ns and 100 s (10 points per decade, isologarithmically). To ensure complete recovery of the dark state, samples were kept in the dark for 120 s before the subsequent recording. For the transient absorption spectra shown in Figure 3h, FTIR samples were used. Spectra were recorded using an Ocean FX array detector (Ocean Optics, Largo, FL) with a spectral resolution of 2.4 nm and integration time of 50 ms. Samples were illuminated using a 50 mW continuous-wave LASER emitting 532-nm light (no. 37028, Edmund Optics, York, UK).

### FTIR spectroscopy

To prepare samples for FTIR, 10 µl of initial protein solution (>20 mg/ml, 100 mM NaCl, 20 mM Tris, 0.05% DDM [pH 8]) was dried stepwise on a BaF_2_ window under a stream of dry air and subsequently rehydrated. Samples were then sealed with a second BaF_2_ window. To ensure constant sample thickness, a 3-µm PTFE spacer was placed between the windows. For deuteration, the protein solution was washed at least five times with deuterium buffer (100 mM NaCl, 20 mM Tris, 0.05% DDM [pD 8]) using Centricon filters and subsequently illuminated using white light to improve deuteration inside the opened channel. FTIR spectra were acquired at 0°C using a Vertex 80v FTIR spectrometer (Bruker Optics, Karlsruhe, Germany) equipped with a liquid nitrogen–cooled MCT detector (Kolmar Technologies, Newburyport, MA). The spectrometer was operated in rapid scan mode with a data acquisition rate of 300 kHz and spectral resolution of 8 cm^−1^. An optical cutoff filter at 1850 cm^−1^ was used in the beamline. After at least 60 min for equilibration, samples were continuously illuminated using a set of green-light LEDs with an emission maximum of 520 nm. Single-turnover illumination was performed using a 10-Hz pulsed Nd:YAG Powerlite 9010 LASER as the pump source for a Horizon II optical parametric oscillator (Continuum, San Jose, CA) set to 530 nm. The pulse width of the setup was approximately 5 ± 2 ns, with an energy output of approximately 60 mJ. The time resolution was 6 ms (achieved by operation in forward-backward mode and splitting of the interferogram).

### RR spectroscopy

RR spectra were acquired with excitation lines of an Ar^+^ (514 nm, 488 nm) and Kr^+^ laser (413 nm) (Coherent, Santa Clara CA). Raman signals were detected in a backscattering configuration (180°) using a confocal LabRamHR spectrometer (Horiba, Villeneuve, France) equipped with a liquid nitrogen–cooled CCD detector. The spectral resolution was approximately 2 cm^−1^. Typical total accumulation time and laser power at the sample were 30 min and 1 mW, respectively. Low-temperature measurements at 80K were carried out with a Linkam cryostat (Linkam Scientific Instruments, Surrey, UK). Samples were inserted into the cell under dimmed red light in order to avoid photoactivation before freezing.

### MD simulations

Classical MD simulations were prepared based on a SWISS homology model^71^ of MerMAID1 on iC++ at pH 8.5 (PDB 6CSN). The iC++ structure was chosen as template as it showed the best combined quality features for structural prediction (best coverage and QMQE [0.61 together with PDB 6CSM], 2^nd^ best QSQE [0.27 vs. 0.28 with PDB 4YZI], and 3^rd^ best identity [32.69% vs. 35.61% with PDB 6EID). The model was prepared using CHARMM-GUI^72^ for the resting state of MerMAID1 with standard protonation for all amino acids. The MerMAID1 monomer was embedded inside a 60 × 60 Å, homogeneous, 1,2-dimyristoyl-sn-glycero-3-phosphocholine bilayer membrane and solvated using a TIP3 water box, adding 10 Å to both the top and bottom of the protein. Systems were simulated under *NPT* conditions using a 2 fs time step, a 303.15 K heat bath, the particle-mesh Ewald method for long-range electrostatics, and the CHARMM36 force field^73^. pK_a_ calculations for all titratable amino acids of MerMAID1 were performed using APBS^74^ in a conformational space of three pH-adapted conformations (PACs) and the Monte Carlo procedure of Karlsberg2+^75,76^ to sample all residues. PACs were created using Karlsberg2+ in a self-consistent cycle including adjustment of protonation patterns of titratable amino acids and salt bridge opening according to pH −10, 7, or 20. To calculate pK_a_ values for MerMAID1 MD frames, only the holoprotein structure was used. Lipids and water molecules were substituted with continuum solvation. Ion channels were predicted using MOLEonline^39^.

### Neuronal recordings and two-photon microscopy

Organotypic slice cultures of rat hippocampus were prepared as described^77^ and transfected by single-cell electroporation^78^ after 14-16 days in vitro (DIV). Plasmids were each diluted to 1 ng/μl in K-gluconate–based solution consisting of (in mM): 135 K-gluconate, 4 MgCl_2_, 4 Na_2_-ATP, 0.4 Na-GTP, 10 Na_2_-phosphocreatine, 3 ascorbate, 0.02 Alexa Fluor 594, and 10 HEPES (pH 7.2). An Axoporator 800A (Molecular Devices) was used to deliver 50 hyperpolarizing pulses (−12 mV, 0.5 ms) at 50 Hz. At DIV 19-21, targeted patch-clamp recordings of transfected neurons were performed under visual guidance using a BX 51WI microscope (Olympus, Shinjuku, Japan) equipped with Dodt-gradient contrast and a Double IPA integrated patch amplifier controlled with SutterPatch software (Sutter Instrument, Novato, CA). Patch pipettes with a tip resistance of 3-4 MΩ were filled with intracellular solution consisting of (in mM): 135 K-gluconate, 4 MgCl_2_, 4 Na_2_-ATP, 0.4 Na-GTP, 10 Na_2_-phosphocreatine, 3 ascorbate, 0.2 EGTA, and 10 HEPES (pH 7.2). Artificial cerebrospinal fluid (ACSF) consisted of (in mM): 135 NaCl, 2.5 KCl, 2 CaCl_2_, 1 MgCl_2_, 10 Na-HEPES, 12.5 D-glucose, 1.25 NaH_2_PO_4_ (pH 7.4). Synaptic currents were blocked with 10 µM CPPene, 10 µM NBQX, and 100 µM picrotoxin (Tocris, Bristol, UK). Measurements were corrected for a liquid junction potential of −14.5 mV. A 16 channel pE-4000 LED light engine (CoolLED, Andover, UK) was used for epifluorescence excitation and delivery of light pulses for optogenetic stimulation (ranging from 385-635 nm). Light intensity was measured in the object plane with a 1918 R power meter equipped with a calibrated 818 ST2 UV/D detector (Newport) and divided by the illuminated field (0.134 mm^2^) of the LUMPLFLN 60XW objective (Olympus).

Neurons in organotypic slice cultures were imaged with two-photon microscopy to characterize their morphology and the subcellular localization of citrine-labeled MerMAID-ChRs. The custom-built two-photon imaging setup was based on an Olympus BX-51WI upright microscope upgraded with a multiphoton imaging package (DF-Scope, Sutter Instrument), and controlled by ScanImage 2017b (Vidrio Technologies, Ashburn, VA). Fluorescence was detected through the objective (NIR Apo 40XW, Nikon, Minato, Japan) using GaAsP-PMTs (Hamamatsu Photonics, Hamamatsu, Japan). A tunable Ti:Sapphire laser (Chameleon Vision-S, Coherent) was set to 810 nm to excite mCerulean, and a high power femtosecond fiber laser (Fidelity-2, Coherent, Santa Clara, CA) was used to excite citrine at 1070 nm.

### Data analysis and statistical methods

Clampfit 10.4 (Molecular Devices) and Origin 2017 (OriginLab, Northampton, MA) were used for analysis of HEK293 electrophysiological recordings. Peak currents were used for analysis of most biophysical properties. The current of the last 50 ms of the illumination period was averaged to determine stationary current amplitude. Reversal potentials were determined based on linear fit of the two data points crossing 0 pA or linear extrapolation from 0 pA most adjacent two data points of a measurement series. Action spectra were normalized to the maximum response and fitted with a three-parametric Weibull function to determine the maximum response wavelength (λ_max_). Kinetic time constants were determined by mono or bi-exponential fits. For displaying reasons electrophysiological recording data points were reduced.

Single turnover UV/vis absorption measurements were averaged over 15 cycles. Primary data analysis was performed using MATLAB R2016b (The MathWorks, Natick, MA) to calculate difference spectra and reconstruct three-dimensional spectra. Glotaran 1.5.1^79,80^ was used for global analysis of the spectral datasets. Time constant values and photointermediate spectra were obtained via global analysis of the data sets. The sequential model explored spectral evolution and produced the EADS, representing the species-associated difference spectra^81^.

Stationary absorption spectra were analyzed using Origin 2017, normalized to maximum absorption at 280 nm or maximum chromophore absorption, smoothed using Savitzki-Golay method using a 10-point window and 5^th^ order polynomial function. Pk_a_-values were determined with a Boltzmann function.

FTIR difference spectra were preprocessed using OPUS 7.5 software (Bruker Optics). FTIR data were analyzed via single value decomposition and rotation procedure and subsequent global fit algorithm implemented in Octave 4.2.^82,83^. Assuming a sequential reaction scheme, a sum of exponential functions was used as the fit model.

RR data was background subtracted with custom written software using a polynomial function.

VMD^84^ and PyMol 2.2.3 were used to analyze and visualize MD simulation results and computed ion permeation pathways.

Neurophysiological data were analyzed and plotted in Igor Pro 8.0.

If not stated otherwise, data was plotted using either MATLAB, GraphPad Prism 7.0 or Origin 2017. Final esthetical adjustments were performed using Adobe Illustrator 2017 (Adobe Systems, San José, CA) or Affinity Designer 1.6 (Serif, Nottingham, UK)

No statistical tests were used to predetermine sample size. Sample sizes were similar to those commonly used in this research field. Repeated experiments always refer to biological replicates performed using at least two batches of transfected cell cultures. Data is given as mean ± standard deviation. Single measurement data and further statistical analysis^85^ are provided in the Supplementary Material. Blinding was not performed to ensure correct assignment of the data to the measured constructs and/or experimental conditions. However, randomization was performed in case of buffer exchange experiments and automated analysis was used whenever possible.

## Data availability

HEK-293 Electrophysiological data is included in the Supplementary Material. Code and data of metagenomic analysis can be obtained from https://github.com/BejaLab/MerMAIDs. Further data or code is available from the corresponding authors upon request. DNA sequences will be deposited on Addgene.

## Acknowledgments

We thank T. Tharmalingam, S. Augustin, M. Reh, M. Meiworm, K. Sauter, und S. Schillemeit for excellent technical assistance. We thank S. P. Tsunoda and H. Kandori for sharing the *Gt*CCR4 plasmid and Itai Sharon for initial *Tara* Ocean assemblies. This work was supported by the Deutsche Forschungsgemeinschaft (DFG; German Research Fundation): SFB 1078 to P.He. (B1, B2), F.B. (B5), and P.Hi. (B6), the Germany’s Excellence Strategy – EXC 2008/1 (UniSysCat) – 390540038 to P.Hi. & P.He., and the priority program SPP 1926 & FOR 2419 to J.S.W. This work was also supported by the European Research Council (ERC): advanced grant (LS1, ERC-2015-AdG) to P.He. and starting grant (LS5, ERC-2016-STG) to J.S.W. P.He. is a Hertie Senior Professor for Neuroscience supported by the Hertie Foundation. O.B. is supported by the Louis and Lyra Richmond Memorial Chair in Life Sciences.

## Author Contributions

J.W., J.O., and P.He. designed the project, with contributions from all authors. J.F.-U., E.P., I.S., O.B., J.W., and J.O. performed computational work. J.O., B.L., S.R-R., P.F., A.S., A.K., and J.V. conducted the experiments, with assistance from M.B. and M.L. Experimental data were analyzed by J.O., J.W., B.L., S.R-R., P.F., A.S., and A.K. and interpreted by all authors. J.W., J.O., and P.He. wrote the manuscript, with contributions from all authors.

