## Supplementary Information for "MerMAIDs: A novel family of metagenomically discovered, marine, anion-conducting and intensely desensitizing channelrhodopsins"

**Oppermann *et al*.**

This file includes:

- Supplementary Discussion
- Supplementary Figures 1-8
- Supplementary Table 1
- Description of additional supplementary files

### Supplementary Discussion

Along with the bands at 1235(-) $cm^{-1}$ and 1199(-) $cm^{-1}$ (Fig 3h) that are typical for depletion of all-*trans* retinal in ChRs[1,2] positive bands arise in the retinal fingerprint region at 1220 $cm^{-1}$ and 1184 $cm^{-1}$ in the fast component spectra that reflect formation of the photoproduct. The band at 1220(+) $cm^{-1}$ was previously observed in mutants of BR and *Cr*ChR2[3,4] and cannot be unambiguously assigned to a distinct mode. The band at 1184(+) $cm^{-1}$, however, indicates formation of 13-*cis* retinal similar to the N intermediate in BR[5,6]. In the slow component spectra, this band is presumably downshifted to 1170 $cm^{-1}$, hinting at an altered chromophore geometry in the desensitized state. Furthermore, the strong negative FTIR band at 1542 $cm^{-1}$, most prominent in the fast component spectra, is due to light-induced changes of $\nu$(C=C) vibrations of the retinal chromophore and correlates with the absorption of the dark state UV-Vis spectrum and the major C=C stretch observed for RR spectra in the dark (Fig. 3e)[7,8]. The amide I region of the fast component furthermore shows a band pattern at 1658(+)/1645(-) $cm^{-1}$ that vanishes upon formation of the desensitized state and presumably reflects small protein backbone rearrangements and an altered retinal C=N configuration during formation of the conducting state (Fig. 3f). In the carboxylic region (>1690 $cm^{-1}$), bands at 1746(+), 1735(-) and 1716(-) $cm^{-1}$ are observed in the fast component spectrum that downshift in $D_2O$ by 3, 3 and 10 $cm^{-1}$, respectively (Fig. 3f and S5f). Therefore, the band pattern at 1746(+)/1735(-) $cm^{-1}$ most likely reflects a hydrogen bond change of a carboxylic residue. Similarly, the band at 1716(-) $cm^{-1}$ is assigned to deprotonation of an aspartic or glutamic acid. In the desensitized state spectra, a strong negative band is observed at 1718 $cm^{-1}$ along with a small positive vibrational mode at 1733 $cm^{-1}$. These bands downshift by 5 and 3 $cm^{-1}$, respectively, after H/D exchange (Fig. 3f and S5f).

### Supplementary Figures

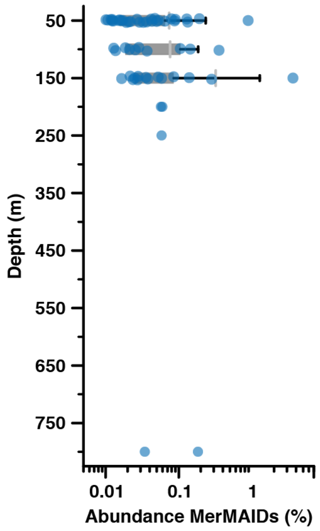


**Fig. S1 | Depth profile of MerMAIDs.** The abundance of MerMAID-like proteins (MerMAID-like/total rhodopsins) was coupled with environmental metadata of the *Tara* Ocean samples to produce the depth profile. Samples were binned into depth groups of ± 25 m. For depth groups with a bin size ≥5, boxes were drawn. Otherwise, only individual data points are shown.

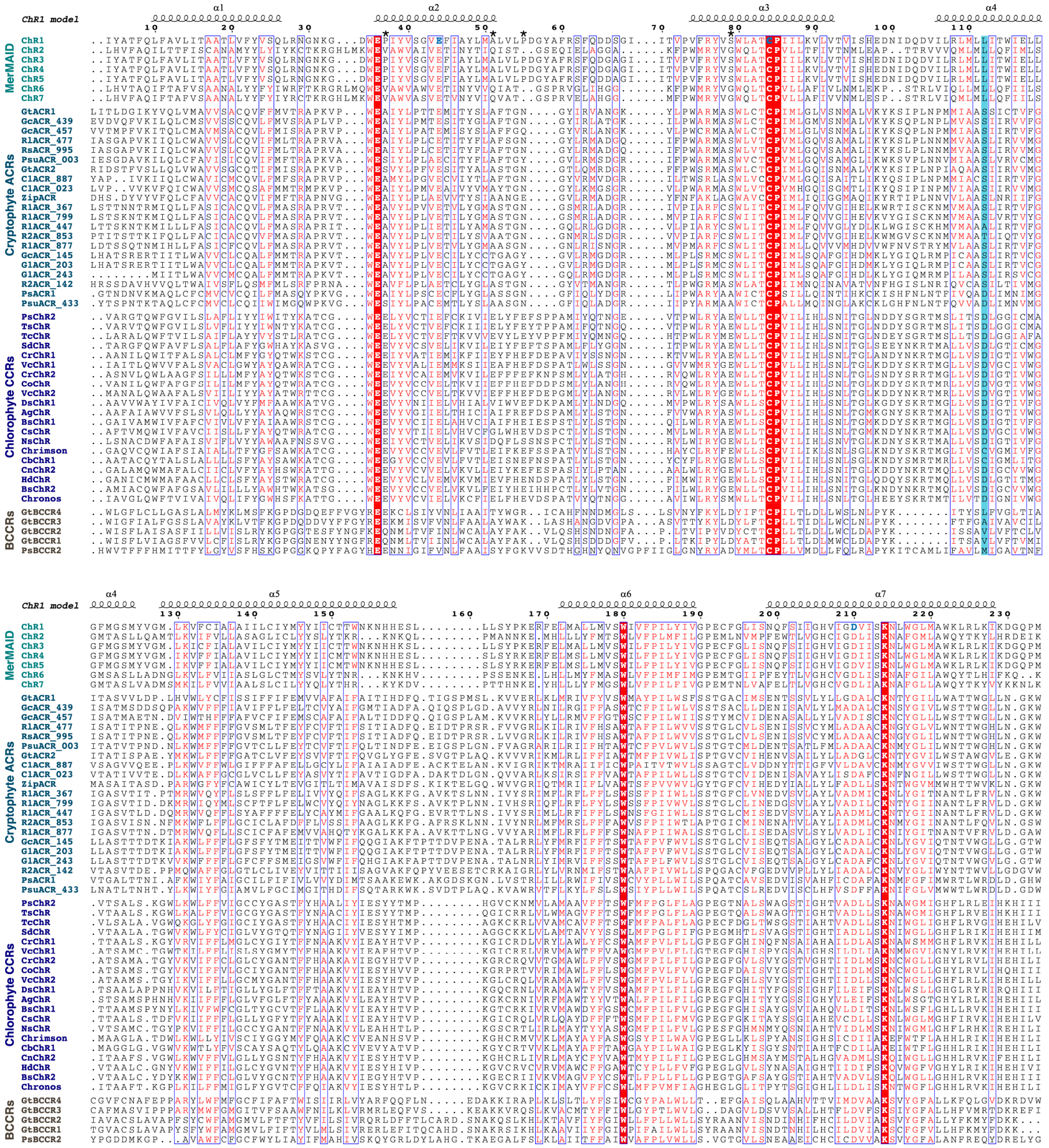


**Fig. S2 | Sequence alignment of selected ChRs.** The α-helices 1-7 of MerMAID1 based on a homology model are indicated above the alignment. Highly conserved residues are highlighted in red in blue boxes. Asterisks indicate positions of typical glutamates in CCRs, neutralized in ACRs. *Cr*ChR2 D156 homolog residues are indicated by light-blue background. Residues, mutated in MerMAID1 and analyzed using electrophysiology, are shown as bold, blue letters. The GenBank IDs and PMIDs of all included ChRs, as well as the full length sequences of MerMAIDs, can be found in Table S2.

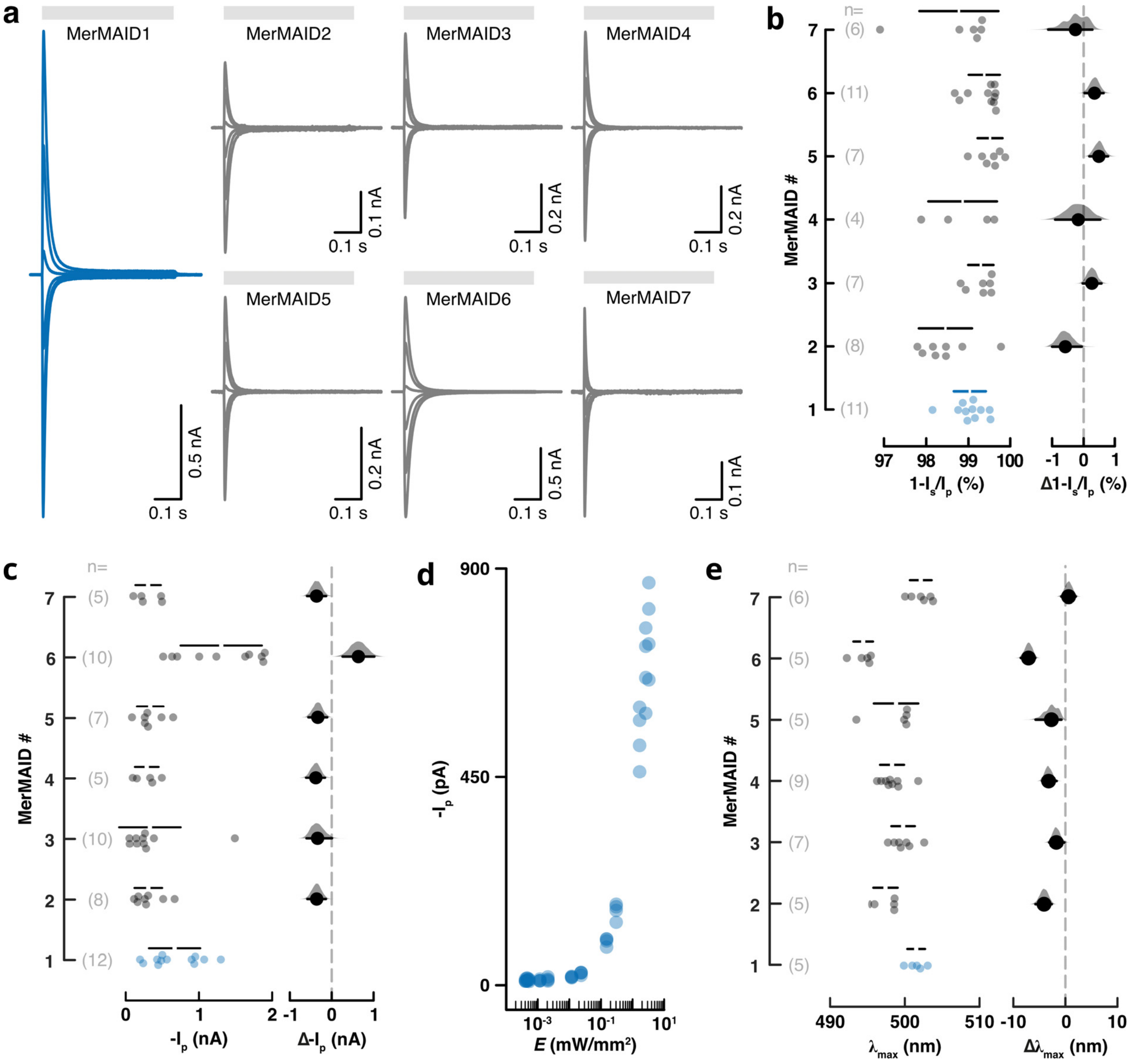


**Fig. S3 | Photocurrent traces and estimation plots for basic MerMAID features. a,** Photocurrents of all MerMAIDs recorded at membrane potentials between -60 mV and +40 mV in steps of 20 mV. Gray bars indicate illumination with 500 nm light. **b, c,** Estimation plots of peak photocurrent decrease (**b**) and amplitudes (**c**) at -60 mV. **d,** Light titration of MerMAID1. Peak photocurrent amplitudes (n=3) are plotted against the light intensity *E*. **e,** Estimation plot of $\lambda_{max}$ for all MerMAIDs. Estimation plots show the mean differences of all MerMAIDs against MerMAID1 on the right as a dot with the respective bootstrap sampling distribution plotted. Error bars indicate 95 % confidence interval. The raw data points are plotted on the left including mean values (white dot) ± standard deviation (black lines). The number of biological replicates measured (n) is denoted.

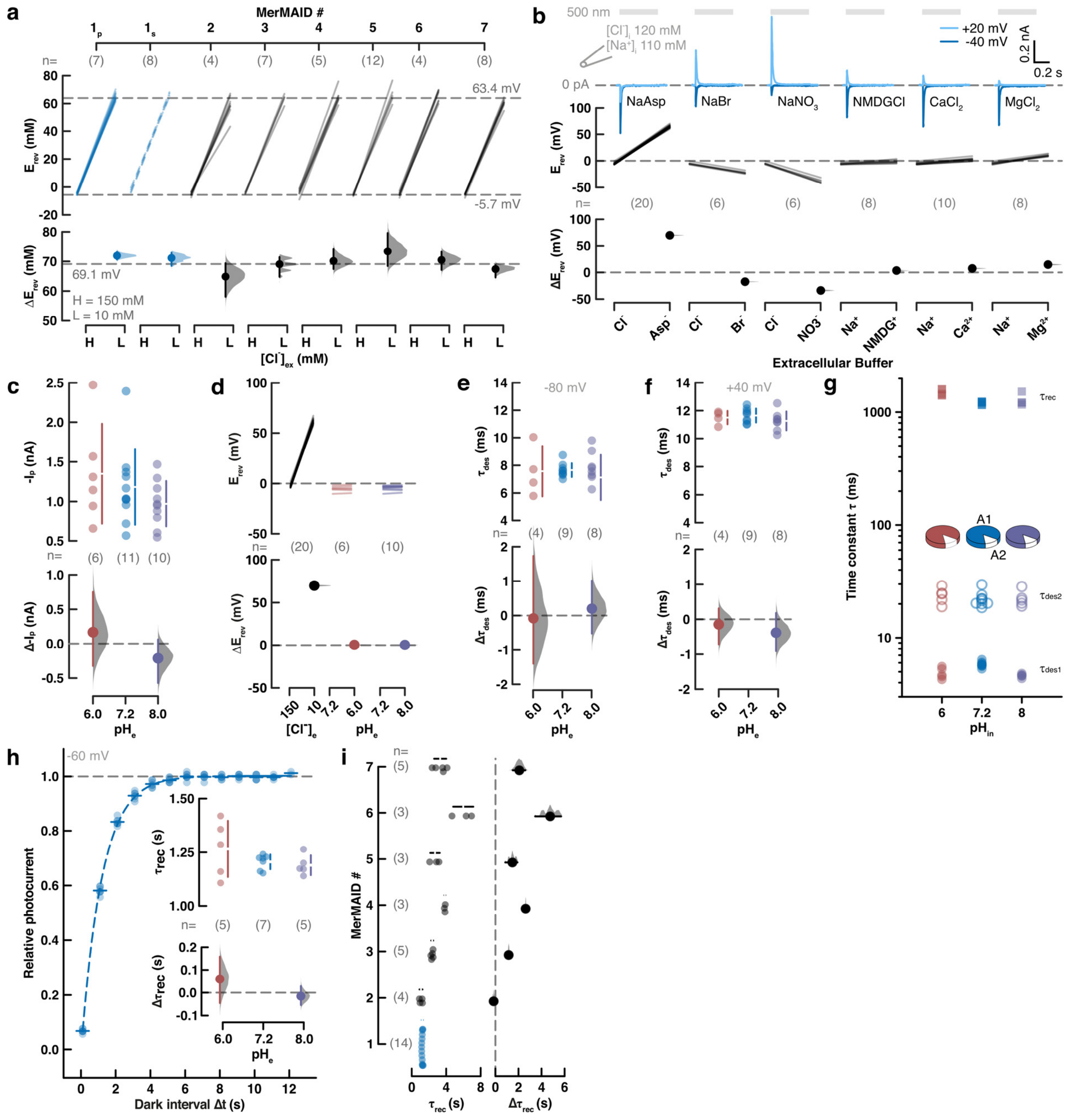

**Fig. S4 | Estimation plots of ion selectivity and kinetic properties of MerMAIDs. a,** (top) Paired reversal potentials ($E_{rev}$) of peak photocurrents at H = 150 mM and L = 10 mM extracellular chloride ($[Cl^-]_{ex}$) of all MerMAIDs and of the stationary photocurrent for MerMAID1. (bottom) Shifts of paired $E_{rev}$ ($\Delta E_{rev}$) upon $[Cl^-]_{ex}$ depletion. Dashed gray lines indicate theoretical Nernst potentials for chloride. **b,** (top) Photocurrents of MerMAID1 recorded at -40 mV or +20 mV with high extracellular concentrations of the indicated ions. Photocurrents were induced with 500 nm light (gray bars). (center) Paired $E_{rev}$ of MerMAID1 peak photocurrents at high external concentrations of the below indicated ions. (bottom) $\Delta E_{rev}$ of MerMAID1 upon exchange of the external buffers. **c, d,** Peak photocurrent amplitudes at -60 mV (**c**) and paired $E_{rev}$ as well as $\Delta E_{rev}$ at indicated external pH ($pH_e$). Paired $E_{rev}$ and $\Delta E_{rev}$ upon reduction of $[Cl^-]_{ex}$ are given as reference (**d**). **e, f,** $pH_e$-dependence of the apparent desensitization time constant ($\tau_{des}$) at -80 mV (e) and +40 mV. **g,** Intracellular pH ($pH_i$)-dependence of the two time constants and their relative amplitudes (pie chart) of the MerMAID1 desensitization and the peak current recovery time constant ($\tau_{rec}$). **h,** Relative recovered MerMAID1 photocurrent with increasing dark intervals between two light pulses. Inset: $pH_e$-dependence of $\tau_{rec}$ of MerMAID1. **i,** Recovery kinetics of all MerMAIDs. In all Estimation plots, the mean difference to the control is shown as a solid dot with the respective bootstrap sampling distribution plotted as a filled gray curve. Ends of error bars indicate 95 % confidence interval. Raw data points are plotted including mean values (white dot) ± standard deviation (black lines). The number of biological replicates measured (n) is denoted.

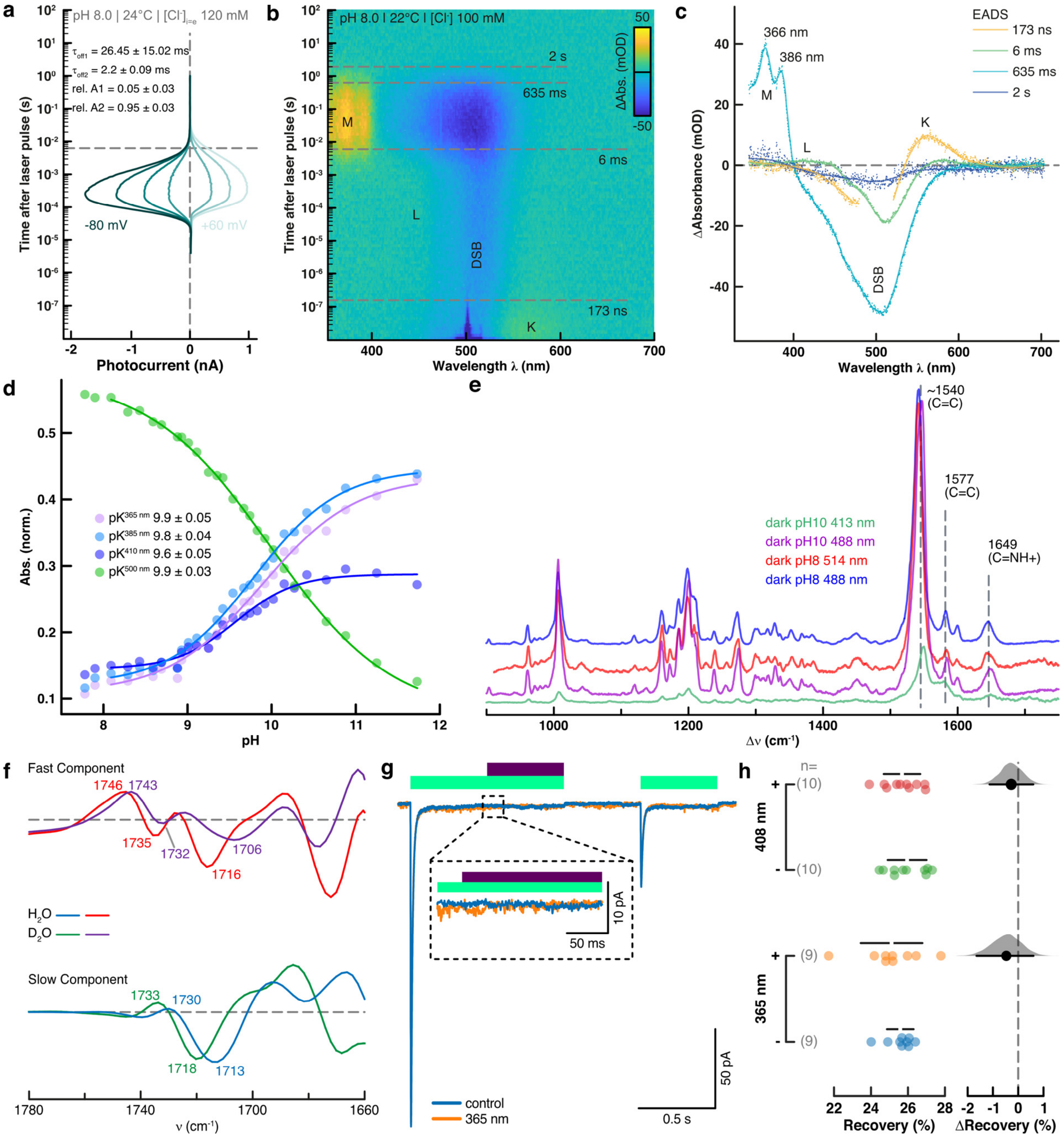

**Fig. S5 | Spectroscopic analysis and M-state illumination of MerMAID-1. a,** Single turnover electrophysiology of MerMAID1 recorded at membrane potentials between -80 mV and +60 mV in steps of -20 mV. The horizontal dashed line indicates L-intermediate decay/M-intermediate rise. Off-Kinetics and relative amplitudes at -80 mV are indicated (n=3). **b,** Reconstructed contour plot from transient absorption spectra of MerMAID1. The photocycle was induced with a 10 ns 500 nm laser-flash. Single letters indicate photointermediates with their respective decay time constants given and indicated by dashed lines. The dark state recovered within 2 s, indicated by the dark-state bleach (DSB) decay. **c,** Evolution associated difference spectra (EADS) resulting from a global fit of the transient absorption spectra are presented as single points where a line is added for visual guidance. Data points saturated from scattered laser light were neglected for performing global analysis of the spectra. Single letters indicate photointermediates. The fine structure peaks of the M photointermediate are indicated as well. **d,** Absorption changes at the indicated wavelengths during changing pH. $pK_a$ values were determined with a Boltzmann function (solid lines). **e,** Resonance Raman spectra of dark-adapted and cryo-trapped MerMAID1 at pH 8/10 recorded with 413 nm, 488 nm, or 514 nm normalized to the peak scattering intensity of the Resonance Raman spectrum recorded with 514 nm. **f,** Kinetically decomposed FTIR light-minus-dark absorption spectra of MerMAID1 in $H_2O$ or $D_2O$ buffer, recorded at 0 °C. **g,** M-intermediate photoreactivity tested by electrophysiology. Photocurrents of MerMAID1 recorded at -60 mV. In a first light pulse, 500 nm light was applied for 1 s. Additionally, the stationary photocurrent was illuminated with 365 nm light for 500 ms. In control experiments, no second wavelength was applied. After a 500 ms dark interval, a second 500 ms 500 nm light pulse was applied to examine potential UV-light induced recovery acceleration. **h,** Estimation plot of the peak photocurrent recovery after additional illumination of the stationary photocurrent with 365 nm or 408 nm. The mean difference to the control is shown as a solid dot with the respective bootstrap sampling distribution plotted as a filled gray curve. Ends of error bars indicate 95 % confidence interval. Raw data points are plotted including mean values (white dot) ± standard deviation (black lines). The number of biological replicates measured (n) is denoted.

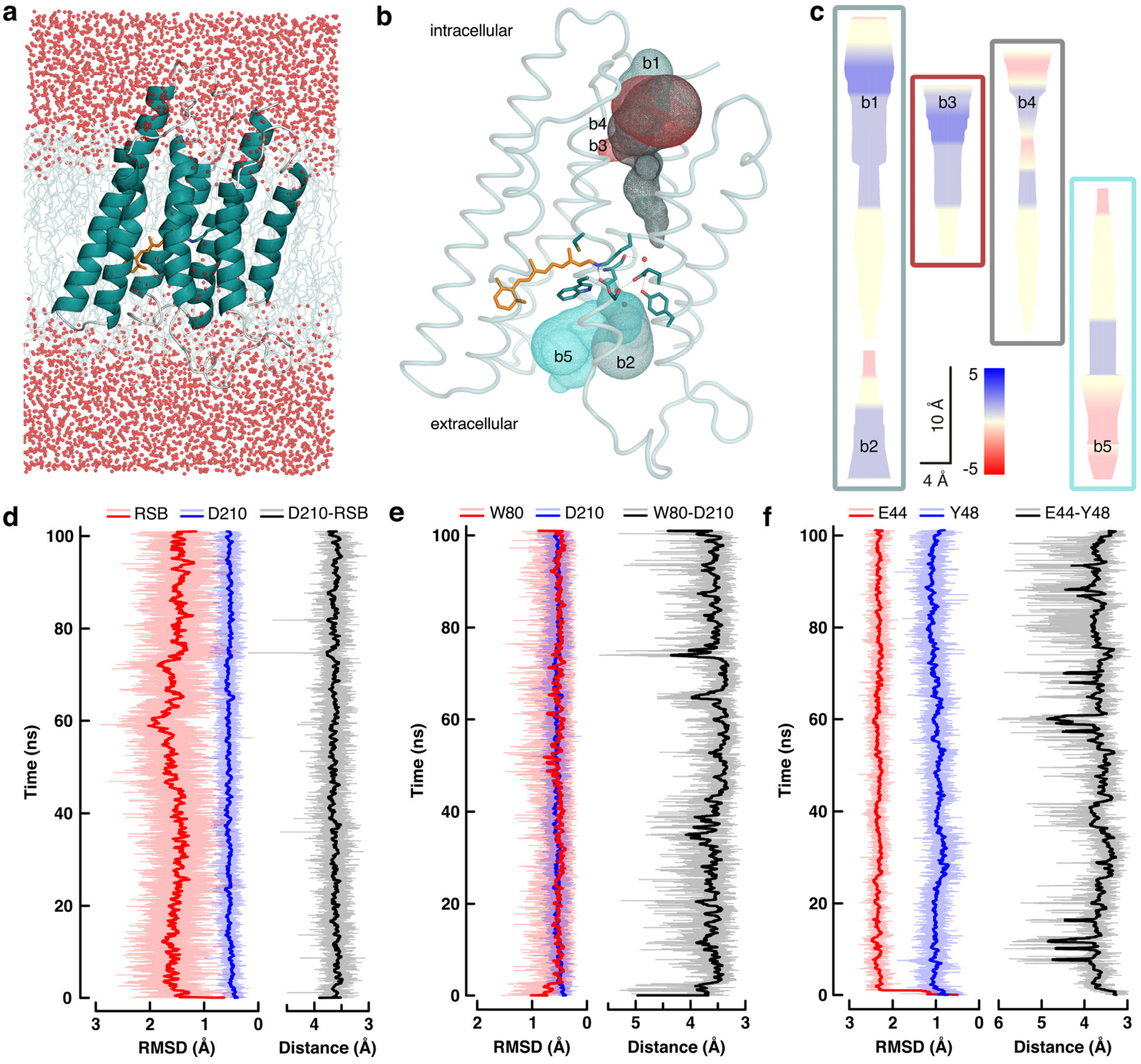


**Fig. S6 | MD simulations and pore prediction of MerMAID1. a,** MerMAID1 homology model embedded in a DPMC bilayer. Red spheres denote water molecules. The protein backbone is represented as cartoon, the retinal as sticks, and lipids as lines. **b,** Dark-state MerMAID1 homology model represented as ribbons. Retinal and active site residues shown as sticks. All predicted ion permeation pathways shown as meshes. Tunnels b1 and b2 were chosen as most likely ion permeation pathway due to most positive surface potential, facilitating chloride transport. Tunnel b4 is predicted to start in the same region as tunnel b3 and then transitions into tunnel b1. **c,** Electrostatic surface potentials and dimensions of the predicted ion permeation pathways. **d-f,** Geometrical sidechain root-mean-square deviation (RMSD) and distance correlations for key residues in the vicinity of the MerMAID1 RSB. Faint graphs show raw data, bold lines are percentile filter smoothened representations.

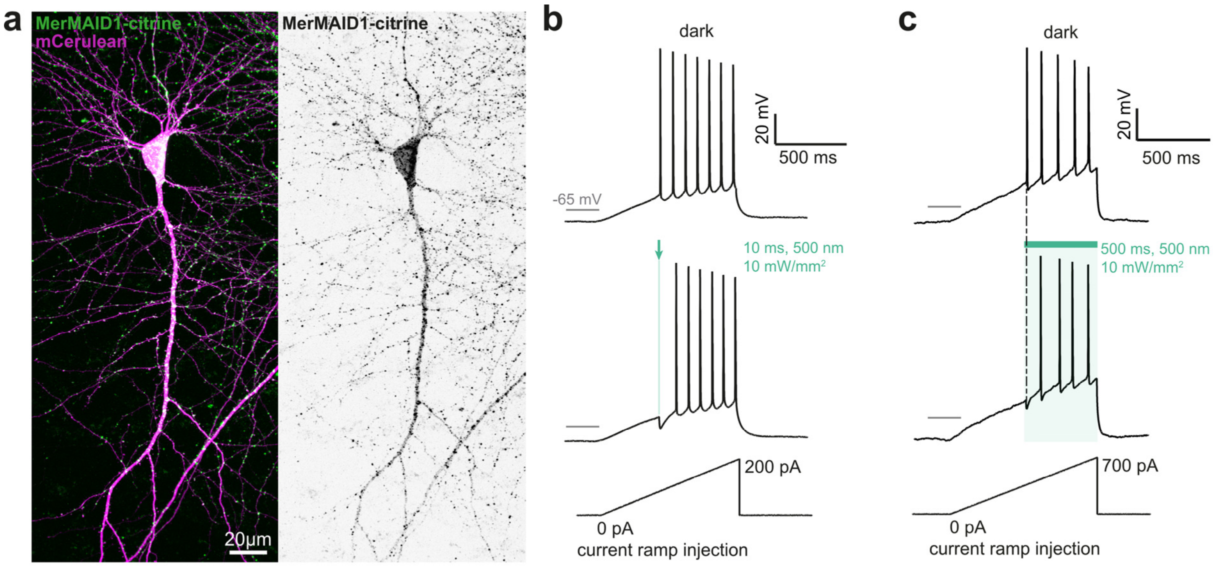


**Fig. S7 | MerMAID1 as an optogenetic silencer in CA1 pyramidal cells in rat organotypic hippocampal slice cultures. a,** CA1 pyramidal neuron expressing MerMAID1-Citrine (green) 5 days after electroporation (stitched maximum intensity projections of two-photon images). mCerulean (magenta) was co-electroporated to visualize neuronal morphology (left). Fluorescence intensity is shown as inverted gray values (right). **b, c,** Voltage traces in response to depolarizing current ramps injected into MerMAID1-expressing CA1 pyramidal cells. Illumination with green light (500 nm, 10 mW/mm$^2$) for a brief (10 ms, **b**) or longer (500 ms, **c**) time blocked single spikes. Light onset preceded action potential onset (measured in the dark condition) by 5 ms.

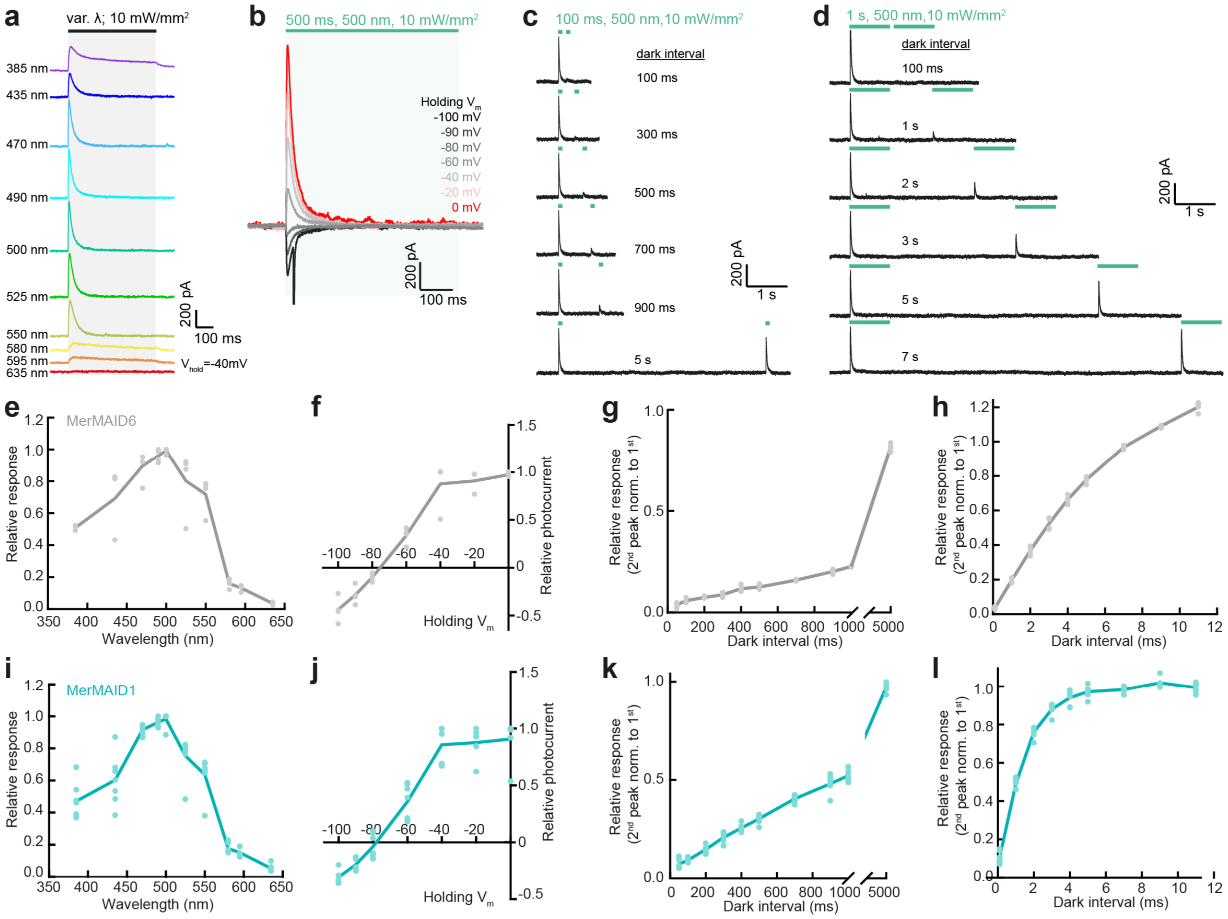


**Fig. S8 | Characterization of spectral activation, ion selectivity and kinetics of MerMAID6 and MerMAID1 in CA1 pyramidal cells in rat organotypic hippocampal slice cultures. a,** Representative photocurrent traces of MerMAID6 elicited with light of various wavelengths for 500 ms with 10 mW/mm$^2$ at a holding potential of -40 mV. **b,** Example photocurrent traces of MerMAID6 elicited with 500 nm light at different holding potentials. **c,** Example photocurrent traces of a double-light pulse experiment at -40 mV to determine the peak current recovery time constant. The dark interval between two light pulses of 100 ms each ranged from 100 ms to 5 s. **d,** Same as (c) but using two light pulses of 1 s each and a dark interval up to 11 s. **e,** Normalized action spectrum of MerMAID6. **f,** Current-voltage relation of the MerMAID6 peak photocurrent. **g,h,** Relative response of MerMAID6 to a second light stimulation after dark intervals of increasing duration. The duration of the two light stimuli was 100 ms in (**g**) and 1 s in (**h**), as shown in example traces (c) and (d), respectively. **i-l,** Same as (e-h) but for MerMAID1. In all graphs, filled circles represent single measurements and solid lines connect mean values.

### Supplementary Tables

**Table S1 | Composition of intra- and extracellular buffers for electrophysiological experiments.** All concentrations are given in mM, LJPs are listed in mV. Asp, Aspartate; EGTA, ethylene glycol tetraacetic acid; HEPES, 4-(2-hydroxyethyl)-1-piperazineethanesulfonic acid; LJP, liquid junction potential; NMG, N-Methyl-D-glucamine

| | | NaCl | KCl | $MgCl_2$ | $CaCl_2$ | CsCl | NaAsp | NMG | HCl | NaBr | $NaNO_3$ | HEPES | EGTA | LJP |
|---|---|---|---|---|---|---|---|---|---|---|---|---|---|---|
| **Intra** | **NaCl** | 110 | 1 | 2 | 2 | 1 | - | - | - | - | - | 10 | 10 | - |
| **Extra** | **NaCl** | 140 | 1 | 2 | 2 | 1 | - | - | - | - | - | 10 | - | 0.6 |
| | **NaAsp** | - | 1 | 2 | 2 | 1 | 140 | - | - | - | - | 10 | - | -12.6 |
| | **NMGCl** | 1 | 1 | 2 | 2 | 1 | - | 140 | 140 | - | - | 10 | - | 6.3 |
| | **CaCl2** | 1 | 1 | 2 | 70 | 1 | - | - | - | - | - | 10 | - | 4.3 |
| | **MgCl2** | 1 | 1 | 70 | 2 | 1 | - | - | - | - | - | 10 | - | 5.0 |
| | **NaBr** | - | 1 | 2 | 2 | 1 | - | - | - | 140 | - | 10 | - | 1.0 |
| | **NaNO3** | - | 1 | 2 | 2 | 1 | - | - | - | - | 140 | 10 | - | -0.3 |

### Description of additional supplementary files

File name: Table S2

Description:

**Table S2 | Overview of ChRs used to generate the phylogenetic tree in Fig. 1a and the sequence alignment in Fig. S2.** For previously described ChRs, AA IDs, GeneBank IDs, and PMIDs are indicated. For MerMAIDs, the SAMEA (SAM, BioSample accession; E, EBI; A, Assay Sample) reads are indicated. The first seven digits are the European Nucelotide Archive (ENA) EMBL sample ID. The second seven digits indicate the contig number of the sample. X denotes two overlapping sequences used to generate the sample

File name: Source Data

Description:

**Source Data | Electrophysiological data and statistics.** The data, used to generate the figures form measurements using MerMAID-expressing HEK293 cells, is given for each figure in a separate sheet. The number of repeated experiments (N), the mean, standard deviation (Std. dev.), standard error of the mean (SEM), and the lower and upper bound of the 95 % confidence interval of the mean are indicated where relevant. The estimation statistics for the main figure panels are given below the raw data and the respective supplementary figure panel is indicated. The estimation statistics comprise the reference and experimental group, mean difference, the low and high endpoints of the bias-corrected and accelerated (BCA) confidence interval (CI; 95 %), and the p-values of a two-sampled independent t-Test and a Mann-Whitney U-Test. All experiments are biological replicates.
